# Mechanistic diversification of *XIST* regulatory network in mammals

**DOI:** 10.1101/689430

**Authors:** Olga Rosspopoff, Christophe Huret, Amanda J. Collier, Miguel Casanova, Peter J. Rugg-Gunn, Jean-François Ouimette, Claire Rougeulle

**Affiliations:** Université de Paris, Epigenetics and Cell Fate, CNRS, F-75013 Paris.; Epigenetics Programme, The Babraham Institute, Cambridge CB22 3AT, UK.; Wellcome Trust – Medical Research Council Cambridge Stem Cell Institute, University of Cambridge, Cambridge CB2 1QR, UK.

**Keywords:** X chromosome inactivation, *XIST*, *JPX*, lncRNA, pluripotency, single-cell RNA-seq, human embryogenesis, evolution, gene regulatory networks

## Abstract

X chromosome inactivation (XCI) is a developmental regulatory process that initiates with remarkable diversity in various mammalian species. Here we addressed the contribution of XCI regulators, most of which are lncRNA genes characterized in the mouse, to this mechanistic diversity. By combining analysis of single-cell RNA-seq data from early human embryogenesis with various functional assays in naïve and primed pluripotent stem cells and in differentiated cells, we demonstrate that *JPX* is a major regulator of *XIST* expression in human and in mouse. However, the underlying mechanisms differ radically between species and require *Jpx* RNA in the mouse and the act of transcription of *JPX* locus in the human. Moreover, biogenesis of *XIST* is affected at different regulatory steps between these species. This study illustrates how diversification of LRGs modes of action during evolution provide opportunities for innovations within constrained gene regulatory networks.

**Graphical abstract:** 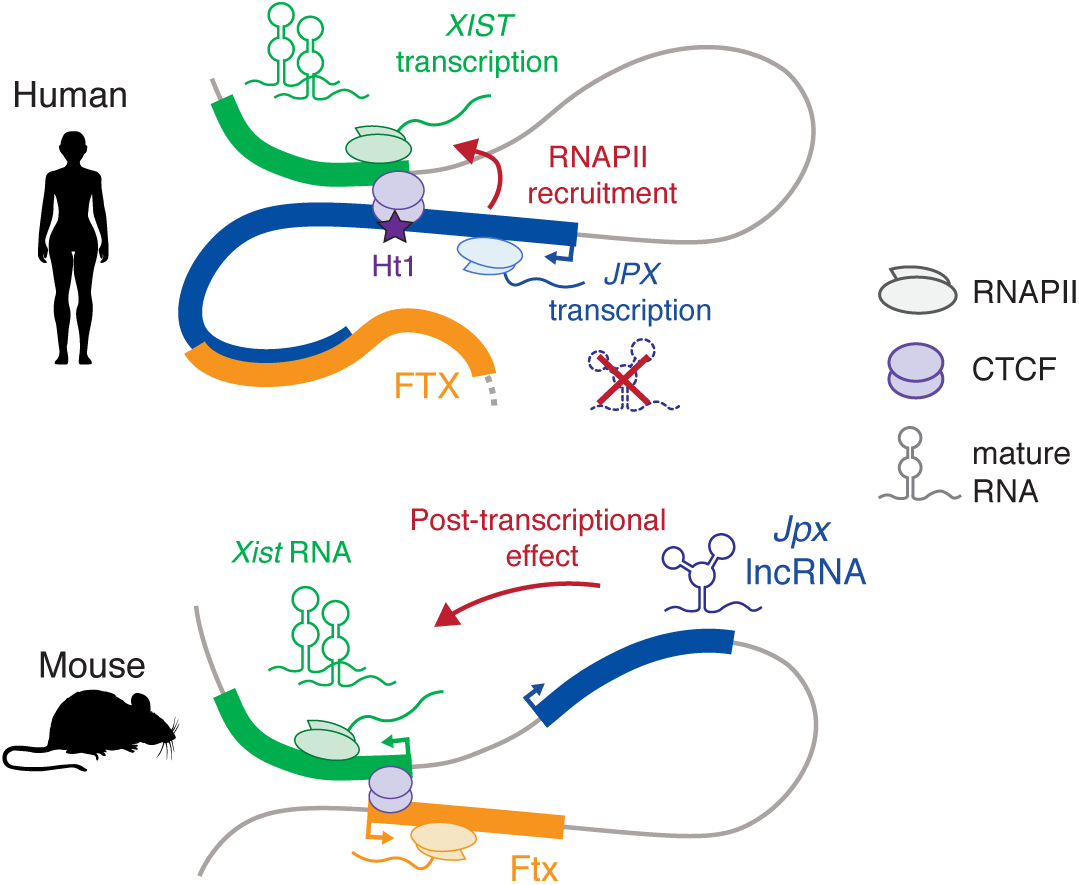

## Introduction

X chromosome inactivation (XCI) is a fundamental epigenetic process that ensures dosage compensation for X-linked genes expression between male and female mammals. X chromosome silencing is triggered early in development by the accumulation of the long non-coding RNA (lncRNA) *XIST*, which acts as a scaffold for multiple protein complexes, involved amongst others in chromatin remodeling, nuclear organization and RNA modification (Furlan and Rougeulle, 2016). The concerted action of these ribonucleoprotein factors results in the conversion of one of the two X chromosomes in females into a compact and transcriptionally silent structure. *XIST* expression has to be tightly controlled in order to ensure female-restricted inactivation of a single X chromosome in a timely manner. However, it remains intriguing that such an essential process follows species-specific routes in which the dynamics of *XIST* expression in early developmental stages differs markedly between mouse and human. While in the mouse *Xist* is restricted to females and to a single X, *XIST* expression in human pre-implantation development transiently initiates in a manner that is independent of the sex and of the number of X-chromosomes. *XIST* accumulation precedes the establishment of proper XCI in human pre-implantation embryos resulting in active *XIST*-coated X-chromosomes (Okamoto et al., 2011). However, the situation eventually homogenizes and *Xist/XIST* RNA coating becomes restricted to a single Xi in both mouse and human post-inactivation (post-XCI) cells (Vallot et al., 2016). These observations raise questions regarding the functional conservation of *XIST* regulatory network across species. We previously identified the transcription factor YY1 as a potent activator of *XIST* in mouse and human (Makhlouf et al., 2014), but the activity of the regulatory elements from the genomic region surrounding *XIST*, known as the X-inactivation center (*XIC*), has never been addressed in species other than the mouse.

Chromosomal rearrangements in mouse and human have allowed the determination of the physical boundaries of the *XIC*, defined as necessary and sufficient to trigger XCI. In addition to *Xist*, other genes were mapped to the mouse *Xic*, including several protein-coding genes (*Slc16a2, Cnbp2, Chic1* and *Rnf12*) and four additional lncRNA genes (*Linx, Tsix, Jpx, and Ftx*). While the order and orientation of *Xic*-linked genes are globally preserved between mouse and human (Chureau et al., 2002; Duret et al., 2006), the human *XIC* underwent a dramatic expansion and is about three times larger compared to its mouse counterpart. Particularly relevant are the lncRNA genes hosted within the *Xic*, which have been linked to *Xist* regulation in the mouse: *Tsix* and *Linx* acts as major repressor of *Xist* expression while *Jpx* and *Ftx* acts as positive regulators. The mechanistic dissection of these *Xic*-linked and other lncRNA loci highlighted that their molecular function is not only mediated by the RNA molecule itself, but may also involve various entities such the act of transcription or key regulatory elements embedded within their locus (Cho et al., 2018; Engreitz et al., 2016; Furlan et al., 2018; Paralkar et al., 2016). For instance, the antisense transcription of the *Tsix* gene over the *Xist* locus contributes to the monoallelic repression of *Xist* (Navarro et al., 2005); *Tsix* transcription is itself controlled by the upstream lncRNA gene, *Linx* (Giorgetti et al., 2014; Nora et al., 2012). *Ftx* transcription has been shown to be essential for *Xist* expression *cis* (Furlan et al., 2018) while *Jpx* was proposed to act through its RNA molecule by binding and titrating away the CTCF protein from the *Xist* promoter (Sun et al., 2013). Therefore, it appears more appropriate to define loci producing lncRNAs as lncRNA genes (LRGs) to better emphasize their mechanistic versatility. The antagonistic action of the *Xic*-linked LRGs is likely facilitated by the spatial segregation of the *Xist*- and *Tsix*-associated regulators in two adjacent and oppositely regulated topologically associated domains (TADs) (Nora et al., 2012). These TADs also delimit internal long-range interactions to ensure contacts between regulatory elements and their target genes; intra-TAD interactions have been described between the *Xist* and *Ftx* LRGs and between *Tsix* promoter and *Linx* LRG (Furlan et al., 2018; Nora et al., 2012).

The molecular players controlling XCI have been largely characterized in the mouse, in part due to the lack of cellular models recapitulating the early stage of *XIST* activation in human. While the differentiation of female mouse embryonic stem cells (ESCs) recapitulates *Xist* upregulation and XCI, no human *ex-vivo* model faithfully reproduces the biallelic upregulation of *XIST* observed during *in vivo* human embryogenesis. Conventional hESCs displays the hallmarks of primed pluripotency including an inactive X-chromosome coated by *XIST* (Vallot et al., 2016). In addition, loss of *XIST* expression may occur spontaneously upon prolonged culture of primed hESCs through a process identified as “XCI erosion”, that involves the ectopic reactivation of a subset of genes from the Xi (Mekhoubad et al., 2012; Vallot et al., 2015). Nevertheless, the post-XCI context in primed hESCs differ from that of differentiated cells and their use as an XCI model has been proven effective in the identification human-specific XCI regulators such as the lncRNA *XACT* (Vallot et al., 2013). It is only recently that methods were developed to reset primed hESCs into the naïve state of pluripotency that matches several features of human pre-implantation embryos, including active X-chromosomes coated by *XIST* (Guo et al., 2017; Sahakyan et al., 2017; Theunissen et al., 2016; Vallot et al., 2016). Indeed, resetting of eroded primed hESCs triggers *XIST* upregulation concomitantly to X-chromosome reactivation (XCR), although *XIST* upregulation is often mono-allelic and restricted to the former Xi (Sahakyan et al., 2017; Vallot et al., 2016). While the resetting of these cells is currently the only method to trigger *XIST* upregulation in human, it has never been used to functionally characterize regulators of *XIST* induction. Altogether, experimental systems are now available to probe the *XIST* regulatory network in human.

From an evolutionary standpoint, one major challenge is that both LRG functionality, if any, and their mechanism of action are hardly predictable based on the DNA sequence alone. The sequence conservation pattern of LRG evolving under functional constraints is therefore difficult to predict. For instance, it is known that syntenic LRG often display strong primary sequence turnover during evolution, even among closely related species (Hezroni et al., 2015; Necsulea et al., 2014; Ulitsky et al., 2011; Washietl et al., 2014), whose impact on LRGs functional conservation is still poorly understood. In rare studies where the functional conservation of lncRNA molecules has been addressed, orthologues display short patches of conserved sequence that are necessary, but not sufficient, for their function (Lin et al., 2014; Ulitsky et al., 2011). While this sharp contrast with the evolutionary stability of protein-coding genes raised controversies on LRGs functionality, experimental investigations have been too limited to provide a definitive understanding of the rules underlying LRGs functional conservation. X-chromosome inactivation provides an interesting experimental paradigm to test this since LRG orthologs are found in the human *XIC* (Romito and Rougeulle, 2011).

In this study, we investigated the regulatory network involved in the initial steps of *XIST* expression in human during early embryogenesis. The analysis of single-cell RNA-seq data from early human embryos designated *JPX* as a potent candidate for *XIST* regulation in human. Using a panel of functional approaches to target various modules of *JPX* LRG, we could show that, while human *JPX* transcripts are dispensable for this process, transcription of the *JPX* locus is essential to sustain *XIST* transcription in post-XCI cells. This process is fostered within a sub-TAD domain that involves RNA polymerase II-mediated 3D interactions. By resetting primed hESCs carrying various deletions of *JPX* promoter region, we demonstrate that this system is suitable to investigate regulators of *XIST* upregulation in human and identify the *JPX* LRG as a major *cis*-regulator of *XIST* transcriptional activation. We also re-addressed the role of *Jpx* RNA, matching cellular models and functional approaches between human and mouse. We could identify that *Jpx* RNA acts as a positive regulator of *Xist* in mouse post-XCI cells and our findings suggest that *Jpx* regulates *Xist* accumulation in a post-transcriptional manner. In addition to identifying of a novel regulator of *XIST* expression in human, these findings provide a striking demonstration of the mechanistic diversification of orthologous LRGs, which sheds new light on the importance of these noncoding elements in defining species-specific regulatory mechanisms within constrained gene regulatory networks.

## RESULTS

### Identification of candidate regulators of *XIST* during early human embryonic development

To examine whether human *XIC*-linked genes are involved in the initial upregulation of *XIST in vivo*, we investigated their expression kinetics during early embryogenesis (Figure 1A), using single-cell RNA-seq datasets obtained from human pre-implantation embryos (Petropoulos et al., 2016; Yan et al., 2013). *XIST* expression initiates between the four- and eight-cell stages (Figure S1A), corresponding to embryonic day 4 (E4, Figure 1B), and increases hereafter, more predominantly in females than in males. While most of *XIC*-linked genes remained lowly expressed throughout pre-implantation development, *RLIM* and *JPX* show the highest levels at the early embryonic days, although they display different expression trajectories (Figure 1C). The expression of the protein-coding gene *RLIM* is the highest at E3 in both male and female embryos and rapidly decrease in the following days, prior to *XIST* induction (Figure S1B). This pattern likely reflects strong maternal inheritance of *RLIM* transcripts, which is consistent with previous observations made in the mouse (Shin et al., 2010). In contrast, low levels of *JPX* could be detected at the 2-4 cells stage, followed by a major burst of expression at the 8-cell stage (Figure S1A) or E4, coinciding with *XIST* initial induction (Figure 1D). Except at the E3 stage, *JPX* was broadly expressed, independently from the sex of the embryos, although *JPX* levels were almost twice in females compared to male embryos, suggesting an early transcription from the two active X-chromosomes (Figure 1D). Using an RPKM threshold to define *XIST* and *JPX* expressing cells in female embryos (Figure 1E), we found that the majority of the cells were expressing either *JPX* or *JPX* and *XIST* concomitantly, with very few *XIST*-only expressing cells at the early embryonic days (E3 and E4, Figure 1F). This pattern suggests that *JPX* activation may shortly precede *XIST* induction and that the two genes become eventually co-expressed in a vast proportion of cells as development progresses. *JPX* and *XIST* expression levels were weakly correlated at early stages of embryogenesis (Figure 1G) and within embryonic lineages (Figure 1H), indicating that *JPX* transcriptional activation, but not the level of its RNA products, may be a prerequisite for its function. These results point toward the *JPX* LRG as a candidate for the regulation of the initial induction of *XIST* expression during pre-implantation development.

**Figure 1.**
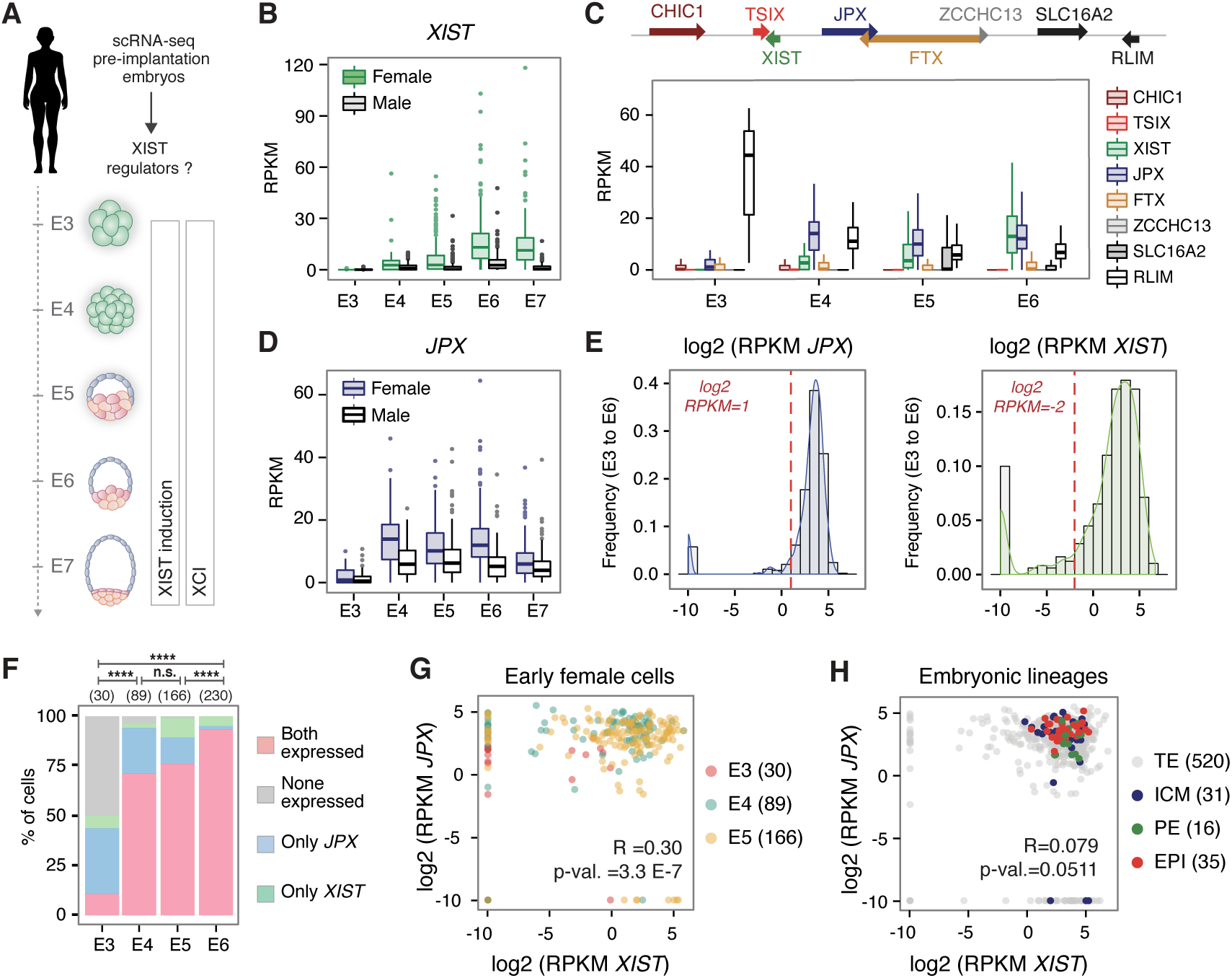
Identification of candidate regulators of *XIST* during early human embryonic development. (A) Single cell RNA-seq data from E3 to E7 pre-implantation embryos (Petropoulos et al., 2016) were used to probe for *XIST* regulators. Also shown is the timing of *XIST* induction along with observed XCI dynamics. (B) *XIST* expression is upregulated in male and female embryos from E4. RPKM: Reads Per Kilobase Million (C) Analysis of single cell expression of *XIC*-linked genes from E3 to E6 reveals that *JPX* induction precedes that of *XIST*. (See also Figures S1A and S1B). (D) *JPX* is expressed with comparable kinetics in male and female embryos. (E) The plots represent the distribution of the log2 RPKM values for *JPX* and *XIST*, in female cells from E3 to E6 stages. Red dashed line represents the cutoff used to define *JPX* and *XIST* expressing cells, on which panel (F) is based. (F) Combined analysis of *JPX* and *XIST* expression in single cells showed that the proportion of cells expressing *JPX* alone decreased during development, while the percentage of cells co-expressing the two genes increased (Chi-square test). n.s., not significant, ***p<0.001; ****p<0.0001. (G-H) *JPX* and *XIST* expression levels were weakly correlated in early stages of embryogenesis and in the different lineages.

### *JPX* RNA is dispensable for *XIST* expression in human

The *JPX* LRG derived from the pseudogenization of the protein-coding gene *USPL* after the divergence of eutherians and marsupials, and evolved concomitantly to *XIST* (Elisaphenko et al., 2008; Hezroni et al., 2017), although the two genes display distinct evolutionary trajectories (Figure 2A). While *XIST* present strong signs of positive selection in both intronic and exonic regions, the *JPX* LRG evolved through a quasi-neutral selection, as illustrated by a conservation score close to zero along the entire locus (Figure 2B). This strong sequence turnover is essentially due to species-specific integration of transposable elements in this region (Chureau et al., 2002; Kolesnikov, 2010), resulting in poor multiple alignment of the homologous region of five eutherian species (Figure 2B). As observed for numerous LRGs (Hezroni et al., 2017; Hezroni et al., 2015; Washietl et al., 2014), signs of purifying selection on the *JPX* gene are concentrated toward the promoter region, including the first exon that contains two highly conserved region of ∼20 nucleotides embedded within the mouse and human transcripts. As the human and mouse genes bear limited sequence identity (Chureau et al., 2002), we examined several features of *JPX* in human, such as its expression pattern and inactivation status in multiple human cell lines. We found that *JPX* expression is not restricted to pre-implantation development but is ubiquitous across a wide range of human tissues (Figure S2A). Similarly to the pre-XCI state, *JPX* transcripts levels appeared consistently higher in females compared to males, suggesting expression from both active and inactive X and, thus, escaping from XCI. To confirm this hypothesis, we performed RNA fluorescence *in situ* hybridization (FISH) to detect simultaneously sites of *JPX* active transcription and of *XIST* RNA accumulation. In both pluripotent and differentiated cellular contexts, *JPX* was expressed in every cell and a pinpoint of transcription was associated *in cis* to *XIST* RNA cloud in about ∼80 to 85% of cells (Figure 2C and S2B), confirming a strong tendency for *JPX* to escape XCI.

**Figure 2:**
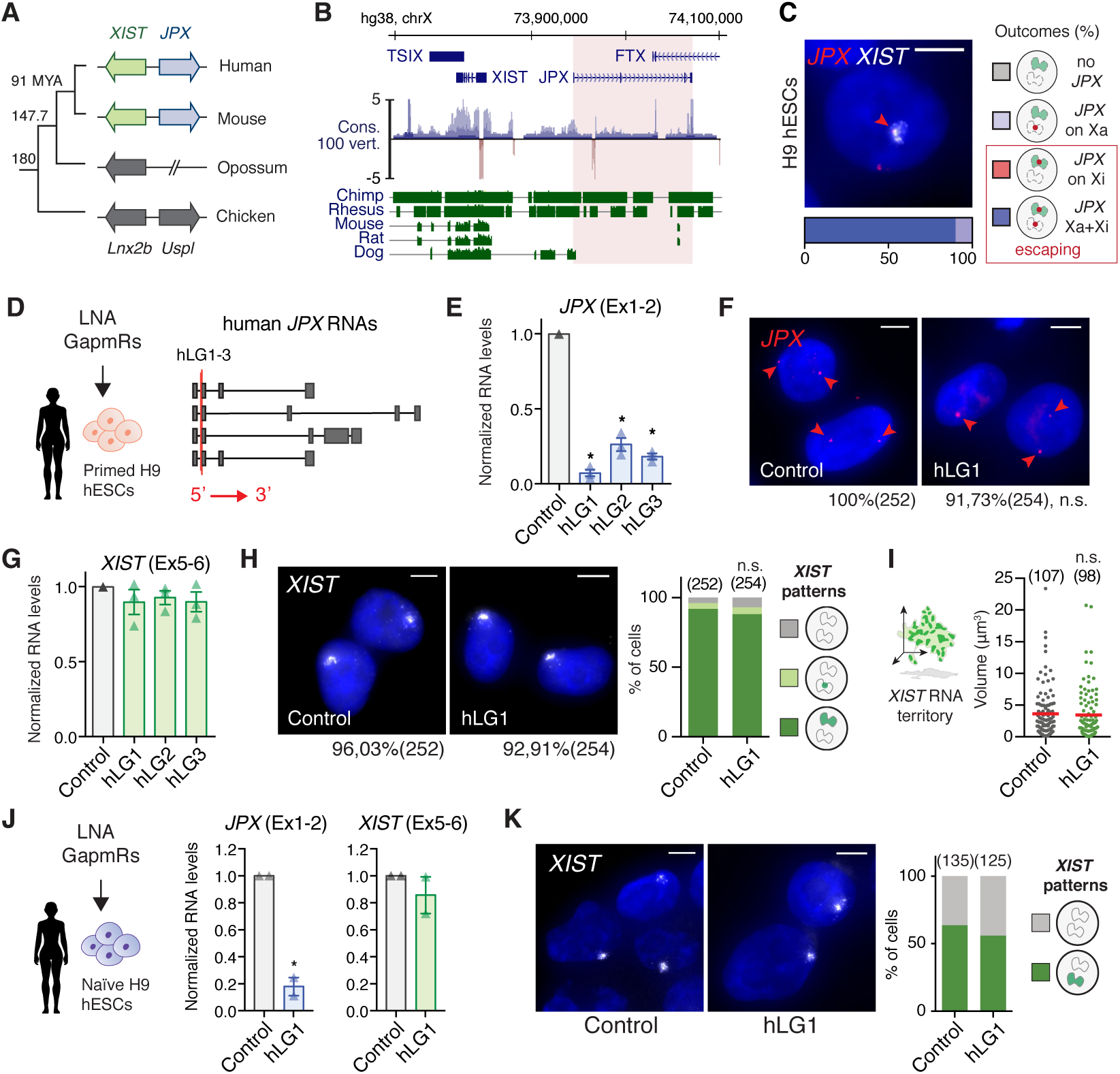
*JPX* RNA is dispensable for *XIST* expression in human. (A) Scheme of *JPX* evolution in vertebrates. Colored arrows represent LRGs and grey arrows their protein-coding ancestors. (B) *JPX* genomic sequence is weakly conserved, as illustrated by the conservation score in 100 vertebrates and alignment of the human genomic region to five other mammalian species. (C) *JPX* escapes XCI in primed H9 hESCs, as assessed by *JPX/XIST* double RNA-FISH (see also Figure S2A and S2B). (D) Scheme of LGs lipofection strategy in primed H9 hESCs and of *JPX* RNA isoforms (red lines: LGs-targeted exons). (E) *JPX* RNA levels are reduced following LG transfection, RT-qPCR, n=3 (see also Figure S2C). (F) LGs targeting human *JPX* RNA did not impact on the number of cells transcribing *JPX* in primed H9 hESCs, RNA-FISH. Percentages indicate *JPX* positive cells, Fischer’s exact test. (G) *XIST* RNA levels are unaffected following *JPX* LG-depletion in primed H9 and WIBR2 hESC (see Figure S2C), RT-qPCR, n=3. (H-I) *JPX* KD did not affect the number of cells expressing *XIST* (Chi-square test) or the volume of *XIST* RNA territory, RNA-FISH, Mann-Whitney test (see also Figure S2D-S2F). (J) In naïve hESCs, *JPX* was efficiently downregulated 48h post-lipofection (unpaired one-tailed t-test, n=2). *JPX* KD did not result in changes in *XIST* RNA levels, RT-qPCR. Expression of pluripotency markers was not affected in these conditions (see Figure S2G), neither was the activity status of the X chromosomes (see Figure S2H). (K) *JPX* KD did not impact on the number of cells expressing *XIST*, RNA-FISH. Scale bars are 5 µm. Error bars represent standard deviation; n.s., not significant; *p<0.05, **p<0.01 and ***p<0.001. Number of counted cells is in brackets.

We next analyzed the function of *JPX* transcripts as they were described as the major functional component in the mouse (Tian et al., 2010). We used an LNA-GapmeR (LGs)-mediated knockdown (KD) strategy (Figure 2D) that have been previously used for the functional analysis of several lncRNAs (Furlan et al., 2018; Leucci et al., 2016; Luo et al., 2016; Tripathi et al., 2010). To address *JPX* function in an embryonic context, we carried out this analysis in non-eroded primed female hESCs, with a percentage of *XIST*-expressing cells above 90% (Vallot et al., 2015). Robust depletion of *JPX* mature transcripts could be achieved in both H9 and WIBR2 lines (Figure 2E and S2C) using three distinct hLGs targeting *JPX* second exon, which is common to all of *JPX* RNA isoforms. As previous studies reported that oligonucleotides containing LNA bases could result in transcriptional inhibition (Beane et al., 2007), we verified that hLGs were not affecting *JPX* nascent transcription by performing *JPX* RNA-FISH (Figure 2F), allowing us to address unambiguously the function of the mature transcripts. In both hESCs lines, *XIST* RNA levels and accumulation within the nuclei remained unaffected by *JPX* KD, as monitored by RT-qPCR (Figure 2G and S2C) and RNA-FISH (Figure 2H-I and S2D). Similar results were obtained in differentiated cells such as fetal fibroblasts (Figure S2E-F), indicating that *JPX* mature RNA is dispensable for *XIST* expression once XCI is established.

Considering that *JPX* is broadly expressed during human pre-implantation development, we hypothesized that *JPX* RNA could function specifically in a pre-XCI context. To test this, we performed *JPX* KD in naïve hESCs that were reset from primed hESCs using recently published methods (Sahakyan et al., 2017; Vallot et al., 2017). These cells display *XIST*-expressing active X-chromosomes and represent the closest *in vitro* model of pre-XCI as observed in pre-implantation human embryos (Guo et al., 2017; Sahakyan et al., 2017; Theunissen et al., 2016; Vallot et al., 2017). *JPX* was efficiently knocked-down in naïve hESCs, without impacting on the expression of naïve-specific markers (Figure S2G). In these conditions, neither *XIST* expression (Figure 2J) nor its pattern of accumulation (Figure 2K) were affected by *JPX* RNA depletion. We also monitored the activity status of the X-chromosomes by monitoring the *ATRX* expression by RNA-FISH as its expression only restricted to fully active X-chromosome (Vallot et al., 2015); *ATRX* transcription remained biallelic in a vast majority of cells suggesting that *JPX* is not involved in XCI *per se* (Figure S2H). Taken together, these results exclude a function of the *JPX* RNA in the expression of *XIST* and XCI in both pre- and post-XCI cells.

### *XIST* expression requires *JPX* transcription

Considering that *JPX* is the closest gene in 5’ to *XIST* (Johnston et al., 2002), we wondered whether *JPX* could be part of the *XIST cis*-regulatory landscape, independently from its RNA transcripts. We characterized long-range interactions surrounding *XIST* promoter region using published Hi-C (Rao et al., 2014) and ChIA-PET datasets (Ji et al., 2016) generated from female cell lines. This revealed a partitioning of the human *XIC* into discrete spatial domains that resembled the organization into topological associated domains (TADs) of the mouse syntenic region (Nora et al., 2012) (Figure 3A). Notably, the boundaries of the *XIST*-associated TAD (∼TAD E) fall within *XIST* locus and upstream of the *RLIM* gene in both species (Figure 3A). Closer inspection of the *XIST*-associated TAD in human (Dowen et al., 2014; Hnisz et al., 2016; Ji et al., 2016) revealed that CTCF-CTCF loops formed an insulated chromatin neighborhood hosting preferential contacts between *XIST* promoter region and the *JPX* gene (Figure 3B). The CTCF loops anchor a region in the vicinity of *XIST* TSS (*XIST*p, +2,9 kb) to two upstream intragenic regions, or hotspots, located within the *JPX* (Ht1, +163 kb) and *FTX* (Ht2, +283 kb) genes. All anchors were co-occupied by CTCF and the cohesin complex, but were not enriched in chromatin marks or proteins associated with active enhancers (Figure S3A). Although a similar organization of the *XIST*-associated TAD was observed on the sole active X chromosome of male fibroblasts (Figure S3B), we found that *XIST/JPX* long-range interactions were associated with RNA polymerase II (RNAPII) only in female datasets (Li et al., 2012), while no peak was reported in that of male (Figure 3B, data not shown). This suggests a female-specific transcriptional association of *XIST* and *JPX* that likely occurs on the Xi (Figure 3B). This hypothesis is further supported by the high number of cells (>80%) co-transcribing *XIST* and *JPX* from the Xi as assessed by RNA-FISH in several cell lines (Figure 2A). These long-range interactions were also detected in CTCF and SMC1 ChIA-PET datasets obtained from female naïve hESCs (Ji et al., 2016), suggesting that these loops occur independently from the XCI status of the cells (Figure S3C).

**Figure 3:**
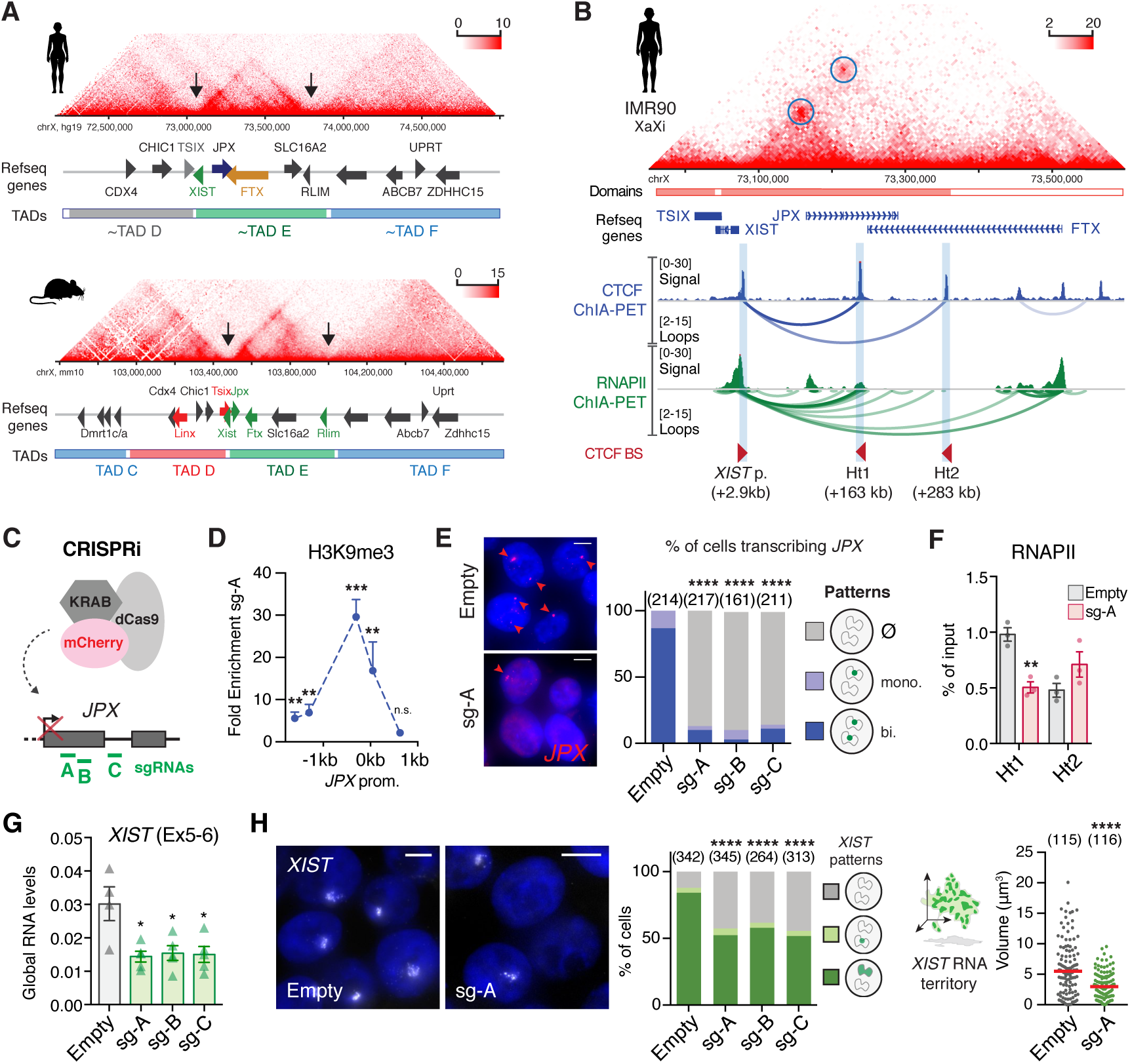
*XIST* expression requires *JPX* transcription. (A) The human *XIC* is partitioned in discrete topological domains (TAD C to F) that are syntenic to that of the mouse *Xic* (TAD D to F); *Xist/XIST* TAD E boundaries are highlighted by black arrows. HiC data from human fetal fibroblasts and mouse ESC (Bonev et al., 2017; Rao et al., 2014). (B) *XIST* and *JPX* interact (Hi-C, IMR90) through CTCF- and RNAPII-mediated loops (ChIA-PET K562). Called loops are highlighted by blue circles. (See also Figure S3A-C). (C-D) Scheme of CRISPRi strategy to inhibit *JPX* transcription in primed hESCs. In this condition, a strong and local enrichment of H3K9me3 could be observed at *JPX* promoter, ChIP-qPCR, n=4. (See also Figure S3D). (E) The number of cells expressing *JPX* was strongly reduced in CRISPRi conditions, with both *JPX* alleles being efficiently silenced one week after lentiviral infection with the guides. Left: Representative images. Right: scoring of *JPX* RNA-FISH signals (Chi-square test). (See also Figure S3E). (F) Inhibition of *JPX* transcription reduced RNAPII availability at the Ht1, ChIP-qPCR, n=3. (G) *XIST* steady state RNA levels were reduced upon inhibition of *JPX* transcription (RT-qPCR, n=4). (See also Figure S3F). (H) *JPX* CRISPRi resulted in a decrease in the number of cells expressing *XIST* (Chi-square test) and on the volume of *XIST* RNA cloud (Mann-Whitney test), RNA-FISH. (See also Figures S3H-K). Error bars represent standard deviation; n.s., not significant; *p<0.05; **p<0.01; ***p<0.001; ****p<0.0001. Unpaired two-tailed t-tests to the empty condition unless stated otherwise. Number of counted cells is in brackets.

As insulated chromatin neighborhoods have been shown to favor the communication between genes and their regulatory elements (Ji et al., 2016; Sun et al., 2019), we hypothesized that *XIST*/*JPX* long-range interactions could provide a structural framework for *JPX* transcription to regulate *XIST* expression. To test this hypothesis, we used a CRISPR inhibition strategy (CRISPRi) (Gilbert et al., 2013) in female primed H9 hESCs, where three guide RNAs were used independently to recruit a catalytically inactive Cas9 fused to a KRAB co-repressor to the *JPX* 5’ region, in order to prevent its transcription (Figure 3C). This system efficiently triggered the local deposition of the H3K9me3 repressive mark on a restricted region surrounding *JPX* transcription start site (TSS) (Figure 3D). As a result, RNAPII recruitment to *JPX* TSS was compromised (Figure S3D) and transcription at the locus was severely impaired; *JPX* remained transcribed in less than ∼15% of the cells as assessed by RNA-FISH (Figure 3E) and *JPX* RNA levels were reduced by 90% (Figure S3E). RNAPII occupancy was also decreased at the Ht1 (Figure 3F), indicating that this strategy efficiently reduced RNAPII processing along the 70 kb of the *JPX* gene. In these conditions, *XIST* steady-state RNA levels were significantly reduced (Figure 3G), as were the percentage of cells with *XIST* accumulation (Figure 3H) and focal enrichment of the H3K27me3 repressive mark (Figure S3F). In addition, the cells that retained *XIST* expression displayed smaller *XIST* RNA cloud compared to the control condition (Figure 3H), indicating that all cells in the population were affected by *JPX* inhibition. As *XIST* repression may result from XCI erosion, we tested whether inhibition of *JPX* transcription could favor this process. For this, we monitored the accumulation of the *XACT* lncRNA and the transcription of the *POLA1* gene from the Xi by RNA-FISH, as the expression of these genes from the Xi are early markers of XCI erosion and precedes the loss of *XIST* expression in H9 hESCs (Vallot et al., 2015). Inhibition of *JPX* transcription did not trigger either *XACT* or *POLA1* transcription from the *XIST*-coated Xi (Figure S3G) and the two genes remained solely expressed from the active X. Moreover, we could not link the reduction of *XIST* expression to perturbation of YY1 binding on *XIST* promoter (Figure S3H), a known regulator of *XIST* expression in human cells (Makhlouf et al., 2014), suggesting that *JPX* acts through YY1-independent mechanisms. We also verified that reduced *XIST* expression did not result from an ectopic deposition of H3K9me3 at *XIST* promoter due to the CRISPRi strategy (Figure S3I). Altogether, our results, which were reproduced in another primed hESCs line (WIBR2, Figure S3J-K), demonstrate that *JPX* transcription is required for proper *XIST* expression.

### *XIST* expression requires a functional *JPX* allele in *cis*

To further explore the contribution of *JPX* transcription to *XIST* regulation, we generated deletions of *JPX* promoter region in primed hESCs using the CRISPR-Cas9 technology. Based on *JPX* promoter features and its expression in H9 cells, we designed guides RNAs to delete a ∼7kb region encompassing the three first exons of *JPX* (Figure S4A). We developed a strategy where hESCs co-transfected with two sgRNAs, each targeting a region upstream and downstream of *JPX* TSS, could be selected based on the expression of a fluorescent gene, GFP and mCherry, respectively (Figure S4B). Therefore, FACS-sorting of double positive cells maximizes the probability to obtain clones with a direct deletion of *JPX* promoter. This approach allowed us to interrogate the contribution of *JPX* transcription to *XIST* regulation in an allele-specific manner, and tease apart *cis*- from *trans*-effects of the LRG (Figure 4A). However, as no SNP could be identified in H9 cells within the deleted region, we systematically performed simultaneous *JPX* and *ATRX* RNA-FISH to determine on which of the two X-chromosomes (Xa or Xi) *JPX* was still transcribed. Using this strategy, we selected two clones carrying heterozygous deletions of *JPX* promoter region for further investigation, in which the deletion occurred either on the active X-chromosome (Δ*JPX*-Xa) or on the inactive X (Δ*JPX*-Xi) (Figure 4B and S4C-D). Interestingly, the two clones displayed different *JPX* RNA levels depending of the deleted allele, with the expression level in Δ*JPX*-Xa clone reaching only ∼15% of that of the WT clone, and ∼80% in the Δ*JPX*-Xi clone (Figure 4C). This is in agreement with *JPX* being predominantly expressed from the Xa in WT cells, as found for most genes that escape XCI (Carrel et al., 1999). Remarkably, we found that *XIST* expression was perturbed exclusively in the Δ*JPX*-Xi clone with RNA; *XIST* RNA levels were reduced by half compared to the WT and Δ*JPX*-Xa clone (Figure 4D) with only ∼55% of the cells displaying *XIST* RNA accumulation (Figure 4E) and H3K27me3 foci (Figure S4E). Moreover, *XIST* RNA territory in the remaining *XIST*-positive cells were significantly smaller in the Δ*JPX*-Xi clone (Figure 4F), indicating that *XIST* expression was also impacted in those cells. Altogether these results show that transcription originating from the *JPX* promoter region is required to sustain *XIST* expression in *cis* in human post-XCI cells. Moreover, the fact that *JPX* RNA levels were the least perturbed in the Δ*JPX*-Xi clone (Figure 4C) confirmed that *JPX* RNA does not control *XIST* expression in human.

**Figure 4:**
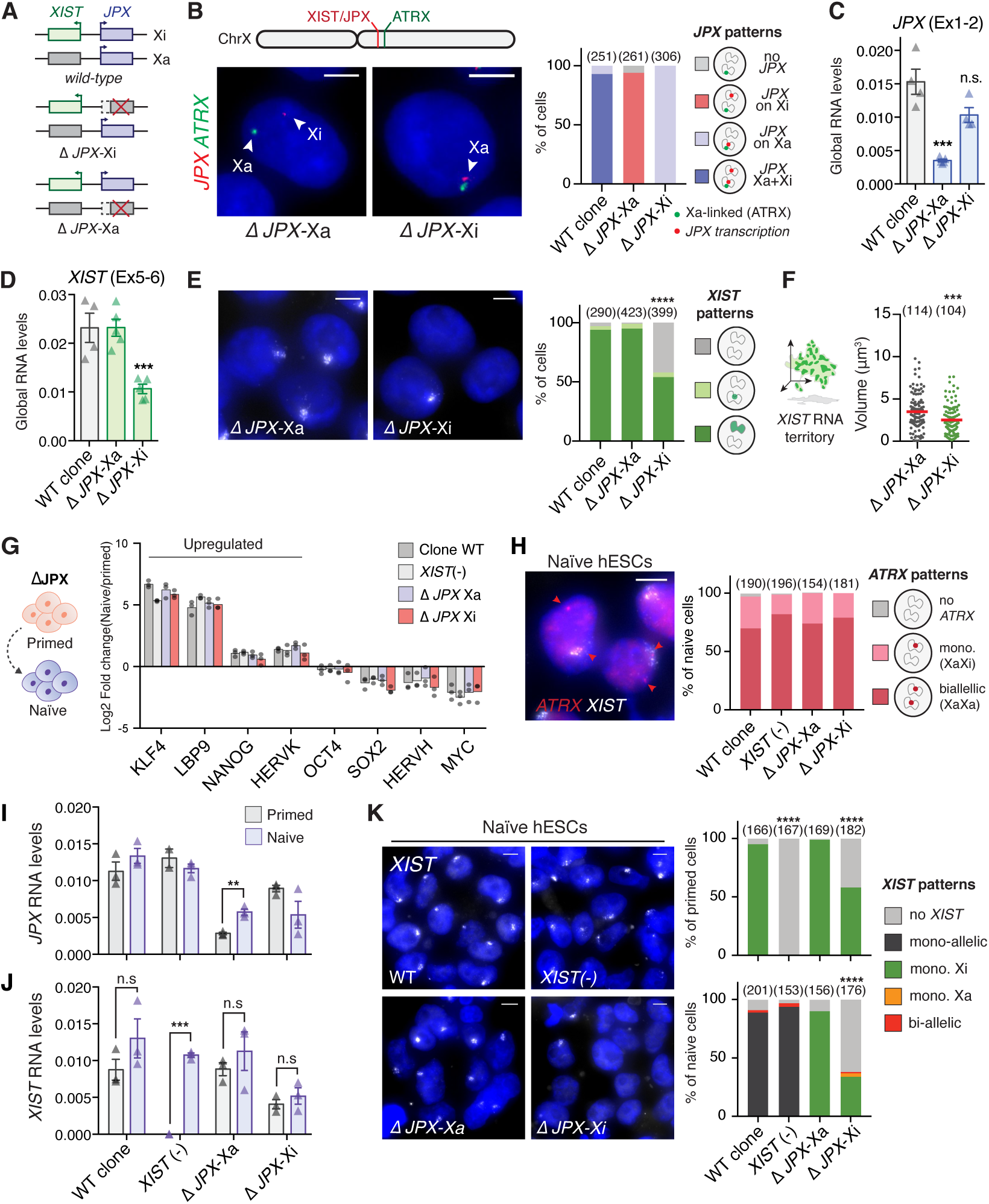
*XIST* expression requires a functional *JPX* allele in *cis*. (A) Schematic representation of WT and *JPX*-deleted hESC clones. (See also Figure S4A and S4B) (B) Determination of the *JPX*-targeted allele by simultaneous *JPX*/*ATRX* RNA-FISH. *ATRX* is transcribed exclusively from the Xa in primed hESCs. (See also Figure S4C and S4D). (C) Δ*JPX*-Xa cells displayed a ∼80% reduction of *JPX* RNA levels compared to WT, while a moderate decrease was observed in Δ*JPX*-Xi cells (∼20%), suggesting an asymmetric expression of *JPX* from the two X chromosomes, RT-qPCR, n=4. (D) *XIST* steady states RNA levels were reduced when *JPX* promoter was deleted in *cis* (*ΔJPX-*Xi), but not in *trans* (*ΔJPX-*Xa), RT-qPCR, n=4. (E-F) In Δ*JPX*-Xi cells, both the number of cells expressing *XIST* (Chi-square test) and the volume of *XIST* RNA cloud (Mann-Whitney test) were reduced, RNA-FISH. (See also Figure S4E). (G) Average log2 fold-change of transcript levels between the naïve and the parental primed hESC clones for a selection of markers, RT-qPCR, n=3. (See also Figure S4F). (H) The Xi was properly reactivated in more than 70% of cells in the different cell lines based on biallelic expression of *ATRX*, RNA-FISH. (See also Figure S4G).

As *JPX* induction seems to precede *XIST* upregulation during human pre-implantation development, we investigated whether its transcription could be important for the *de novo* induction of *XIST* expression. To do so, we proceeded to the chemical resetting of the primed Δ*JPX* and WT lines into the naïve state of pluripotency (Guo et al., 2017). As a control, we also converted fully *XIST*-negative eroded primed hESCs to ensure that the resetting process could efficiently trigger *XIST* upregulation and X-chromosome reactivation (XCR) as observed with this method (Guo et al., 2017) and others (Sahakyan et al., 2017; Theunissen et al., 2016; Vallot et al., 2017). After 7 passages (∼45 days) in naïve culture medium, all cell lines displayed dome-shaped colonies (Figure S4F) and proper induction of key naïve-specific pluripotency markers (Figure 4G) such as the transcription factors KLF4 and LBP9 (Takashima et al., 2014), indicating an efficient transition to the naïve-like state. XCR was observed to same extent in all cell lines, as inferred from the bi-allelic transcription of *ATRX* (∼75-85%, Figure 4H), which confirmed that XCR occurred independently from the *XIST*-expressing status of the parental primed hESCs (Sahakyan et al., 2017). XCR could also be detected at the level of *XIC-*linked genes, notably by the biallelic transcription of the Xa-specific *FTX* gene (Figure S4G) and for *JPX*, which is more expressed in naïve Δ*JPX*-Xa cells compared to primed (Figure 4I). We found that *XIST* was strongly reactivated upon the resetting of the eroded cells (Figure 4J), with the proportion of naïve *XIST*-expressing cells reaching about 95%, as assessed by RNA-FISH (Figure 4K). In agreement with previous studies, *XIST* accumulation remained mostly monoallelic and likely confined to the former Xi, with less than 3% of XaXa cells displaying biallelic *XIST* RNA clouds. Resetting of *XIST*-expressing cells (WT and Δ*JPX*-Xa clones) did not increase the proportion of biallelic *XIST* RNA clouds and *XIST* RNA clouds remained associated to the former Xi (Sahakyan et al., 2017). By contrast, *XIST* reactivation could not be observed in cells where *JPX* is deleted on the Xi allele, in which *XIST* expression remained low and restricted to an even lower percentage of cells, compared to primed conditions (38% naïve vs. 58% primed, Figure 4K). Deletion of the *JPX* promoter region thus prevents *de novo XIST* upregulation in *cis* during conversion of primed to naïve human ESCs. These analyses altogether suggest that *JPX* transcription is required for proper *XIST* upregulation in *cis* during early human pre-implantation development.

### *Jpx* RNA regulates *XIST* expression in mouse post-XCI cells

Previous studies have shown that the deletion of a single allele of the *Jpx* gene or shRNA-mediated knockdown of *Jpx* is sufficient to prevent *Xist* upregulation (Carmona et al., 2018; Tian et al., 2010), with *Jpx* acting through RNA-based mechanisms, both in *cis* and in *trans*. Considering the discrepancy with the results we obtained in human, we decided to revisit the function of *Jpx* RNA in the mouse, by rigorously matching both our experimental approaches and the cellular models. We performed LGs-mediated KD experiment in mouse embryonic fibroblasts (pMEFs) and murine epiblast-derived stem cells (EpiSCs). Murine EpiSCs share similarities with primed hESCs in terms of transcriptional signatures, signaling pathways and XCI status (Brons et al., 2007; Tesar et al., 2007), while pMEFs parallel the primary fibroblasts of fetal origin used in this study (Figure 2C-H and S2F-G). Both cell types display one inactive X-chromosome coated by *Xist* and express *Jpx*. First, we investigated mouse *Jpx* RNA function in primary mouse embryonic fibroblasts (pMEFs) derived from 13.5 days post-coitum mouse embryos using three distinct LNA Gapmers targeting different exons, to minimize the probability of random off-target effects (Stojic et al., 2018) (Figure 5A). This approach was efficient in depleting *Jpx* RNA (Figure 5B), and the strongest effect was obtained with the mLG1, which is expected to target all *Jpx* RNA isoforms. The three LGs induced a decrease in spliced *Xist* RNA levels (Figure 5C) that correlated with the extent of *Jpx* RNA depletion (Pearson correlation = 0.96, p-val. = 0.035), suggesting a dose-dependent effect of *Jpx* RNA. RNA-FISH analyses revealed that most of the cells were affected by *Jpx* KD, as both the percentage of *Xist* positive cells and the volume of the remaining *Xist* RNA clouds were reduced (Figure 5D). As previously, we verified that the observed effects of *Jpx* mLGs were not due to transcriptional inhibition through quantification of *Jpx* nascent transcripts by RNA-FISH and after ethynyl uridine (EU) incorporation followed by pull-down (Figure S5A-B). We could thus conclude that the observed *Xist* downregulation can be attributed to the depletion of *Jpx* mature transcripts only, and not to alterations of *Jpx* ongoing transcription. Similarly to pMEFs, depletion of *Jpx* RNA in EpiSCs led to a decrease in *Xist* RNA levels (Figure 5E), without ectopic expression of *Xist* negative regulators in the mouse, namely its antisense *Tsix* (Figure 5F) or the pluripotency factors REX1 and KLF4 (Figure 5G) (Navarro et al., 2010). These data demonstrate the contribution of mouse *Jpx* RNA to the maintenance of *Xist* expression in post-XCI cells and confirm the role of *Jpx* as a potent regulator of *Xist*.

**Figure 5:**
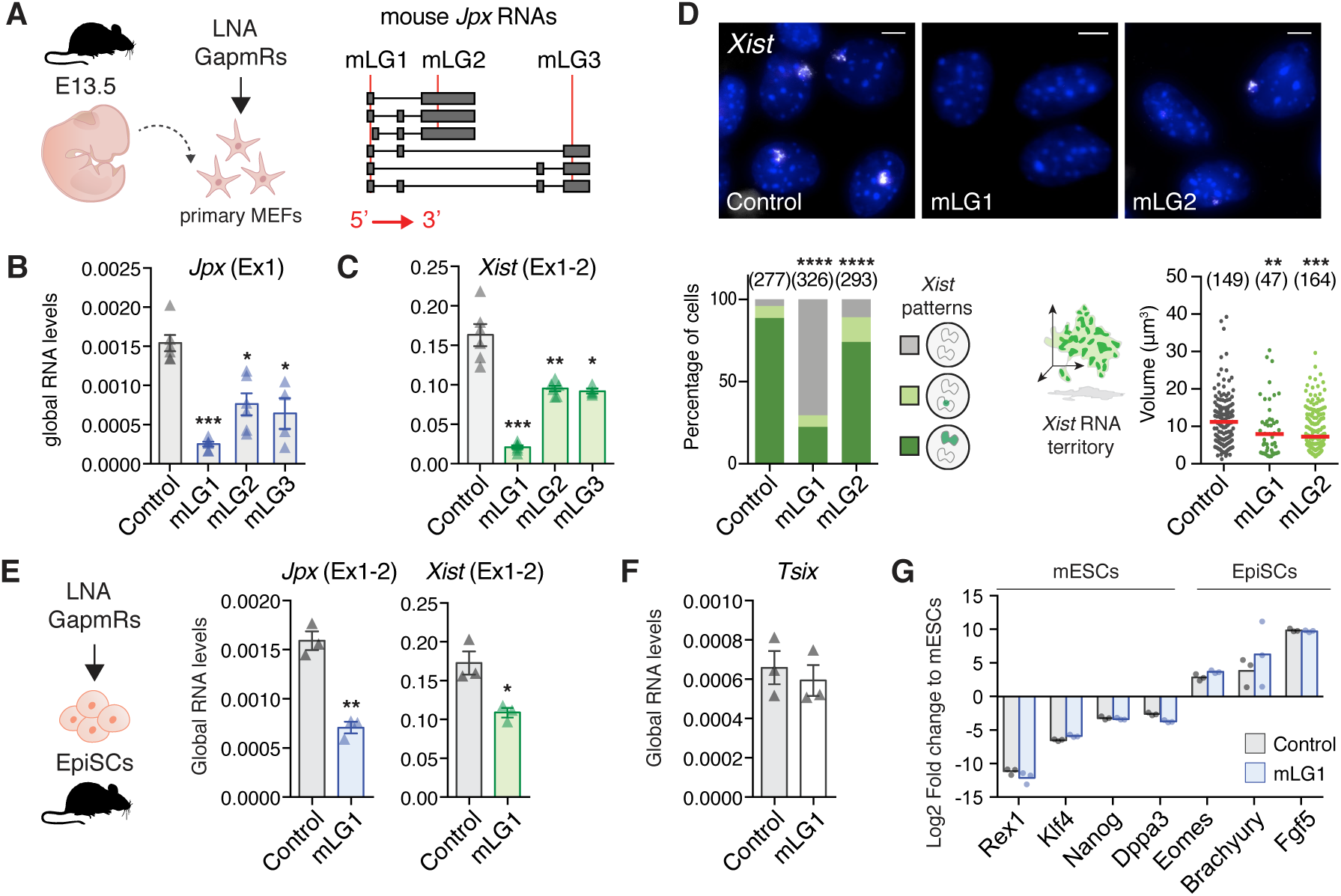
*Jpx* RNA regulates *XIST* expression in mouse post-XCI cells. (A) Schematic representation of LNA GapmeRs (LG) lipofection in primary MEFs; LGs-targeted regions (red lines) are indicated on *Jpx* RNA isoforms. (B-C) LG-transfected pMEFs showed reduced *Jpx* and *Xist* RNA levels, RT-qPCR; n=4. (D) *Jpx* KD reduced the number of *Xist* expressing cells (Chi-square test) and the volume of *Xist* RNA cloud (Mann-Whitney test), RNA-FISH. Red bars: median. (E) In EpiSCs, *Jpx* KD led to a decrease in *Xist* RNA levels, RT-qPCR, n=3. (F) *Tsix* is not re-expressed in EpiSC transfected with *Jpx*-targeting LGs. (G) Log2 expression fold change for a selection of markers (Brons et al., 2007; Tesar et al., 2007) in EpiSC transfected with control or *Jpx*-targeting LG, normalized to expression in mESC, RT-qPCR, n=3. Error bars represent standard deviation; n.s., not significant; *p<0.05; **p<0.01; ***p<0.001; ****p<0.0001. Unpaired two-tailed t-tests to the control LG unless stated otherwise. See also Figure S5.

### Mechanisms of *XIST* regulation by *JPX* have diversified during evolution

To further decipher the mechanisms underlying the different molecular function of *JPX*, we investigated which step of *Xist*/*XIST* biogenesis was under the control of the *Jpx*/*JPX* LRG in mouse and human. Since previous work reported that *Jpx* RNA could activate *Xist* by evicting the CTCF protein from its TSS (Sun et al., 2013), we investigated CTCF binding profile across the *Xist* promoter upon *Jpx* KD in pMEFs. Binding of CTCF to a position ∼1kb upstream of *Xist* TSS was significantly increased upon *Jpx* KD (Figure 6A), an effect we also observed on the imprinting control region of H19 (Figure S6A). Nevertheless, we probed the impact of this change of CTCF binding on *Xist* transcription by measuring the level of *Xist* premature transcripts. Quite unexpectedly, we could not detect changes in *Xist* premature transcript, suggesting that transcription was unaffected by *Jpx* KD (Figure 6B). To confirm this, we performed nascent RNAs pulldown (Figure 6C) and single-cell level by RNA-FISH using stranded oligo-probes detecting *Xist* first intron (Figure 6D). While these approaches were suitable to detect *Xist* transcriptional changes upon *Yy1* KD, a known regulator of *Xist* transcription (Figure 6C-D and Figure S6B-E) (Makhlouf et al., 2014), we could not detect any transcriptional deregulation upon *Jpx* KD. Altogether, these data demonstrate that *Jpx* RNA acts downstream of *Xist* transcription and is required for proper *Xist* RNA accumulation.

**Figure 6:**
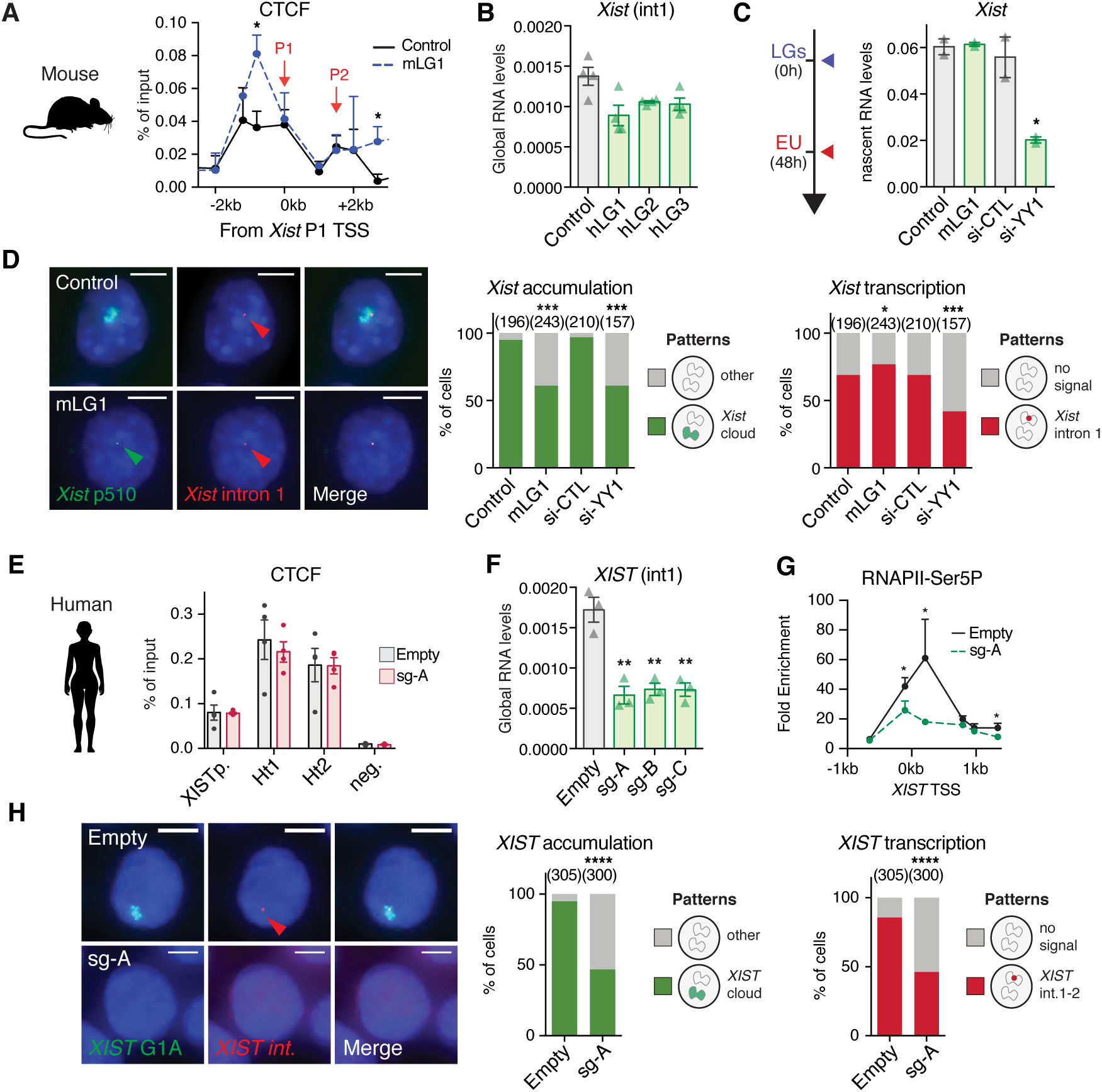
Mechanisms of *XIST* regulation by *JPX* have diversified during evolution. (A) In the mouse, CTCF binding to *Xist* proximal promoter region is increased upon *Jpx* KD in pMEF, ChIP-qPCR, n=3. (see also Figure S6A) (B) KD of *Jpx* RNA did not impact on *Xist* premature transcript levels, intronic RT-qPCR, n=3 (See also Figure S6B-D). (C) *Xist* transcription was affected by *Yy1* KD, but not by *Jpx* KD, in pMEFs when quantified after pulldown of EU-labelled nascent transcripts, RT-qPCR, n=3. (D) *Jpx* KD did not affect *Xist* ongoing transcription (intron 1 stranded oligo-FISH probes) but only on its accumulation (p510 probe) (Fisher’s exact test). *Yy1* KD affects both transcription and accumulation of *Xist*. (See also Figure S6E) (E) In human, *JPX* CRISPRi did not affect CTCF enrichment at *XIST* promoter or at both interaction hotspots, ChIP-qPCR, n=3. (F) *XIST* premature transcripts levels are reduced following *JPX* CRISPRi, intronic RT-qPCR, n=3. (G) Inhibition of *JPX* transcription prevented RNAPII (CTD-phospho-Serine5) recruitment at *XIST* promoter, ChIP-qPCR, n=4. (H) The number of cells with *XIST* RNA accumulation (G1A probe) and transcription (intronic stranded oligo-FISH probes) is reduced following *JPX* CRISPRi. Error bars represent standard deviation; *p<0.05; **p<0.01; ***p<0.001; ****p<0.0001. Unpaired two-tailed t-tests to control condition unless stated otherwise.

In striking contrast, *XIST* unspliced RNA levels were strongly decreased following inhibition of human *JPX* transcription, independently of CTCF binding changes (Figure 6E-F). This was further supported by the observed reduction of phospho-Ser5 RNAPII recruitment at *XIST* promoter (Figure 5c). Severe impairment of *XIST* ongoing transcription following *JPX* transcriptional inhibition was also evident at the single-cell level, when stranded oligo-FISH probes were used to detect human *XIST* intronic regions (Figure 6H). This suggests a transcriptional crosstalk between *JPX* and *XIST* in human, where ongoing transcription across the *JPX* locus would favor the recruitment of the transcription machinery at *XIST*, possibly through local 3D interaction and chromosomal looping. Our results therefore demonstrate that not only the functional module of *Jpx/JPX* differs between human and mouse, but also their mode of action to regulate *Xist/XIST*.

## DISCUSSION

Here, we interrogated the *XIST* regulatory network in human early development, and the extent to which regulatory networks essentially based on LRGs operate similarly in different species. Through unbiased analysis of expression dynamics of *XIC*-linked genes, we identified *JPX* as the best candidate for promoting *XIST* activation, and through functional investigation in various human contexts, we demonstrated a major and ubiquitous role for *JPX* in *XIST* expression. Doing so, we provided the first evidence that resetting human primed to naïve pluripotent stem cell may constitute a system of choice to study the regulatory network at stake for post-fertilization *XIST* activation in human. We further determined that transcription across *JPX* is important in this process and that *JPX* RNA is not involved in *XIST* regulation in human cells. Furthermore, *JPX* acts in *cis* to promote *XIST* transcription, while we show that *Jpx* RNA acts downstream of *Xist* transcription on *Xist* metabolism in the mouse. The choice of cellular models was critical for the comparative analysis of *JPX/Jpx* mode of action in human and mouse. Indeed the early steps of XCI differs markedly in these species, with initial *XIST* up-regulation being uncoupled from XCI in human, and, more importantly, not currently recapitulated in any *ex vivo* model. We therefore chose to focus most of our investigation on the maintenance of *XIST* expression, for which comparable cellular systems were available in the two species studied. The reproducibility of the results in two post-XCI contexts, primed pluripotent stem cells and fetal fibroblasts, strengthens our conclusions and provide the first evidence that an LRG from the *XIC* plays similar function in different mammalian species but operates *via* different mechanisms. *Jpx*/*JPX* therefore stands as a key component of the *Xist*/*XIST* regulatory network in mouse and human.

Addressing the functional conservation of LRGs is a challenge given their fast evolutionary rate. Overexpression of human *JPX* RNA in *trans* was recently shown to complement heterozygous deletion of mouse *Jpx* during the establishment of XCI, suggesting that the human RNA might be functional in an ectopic context (Karner et al., 2019). Such rescue experiments using orthologous LRGs, as opposed to our strategy to tackle the role of mouse and human *Jpx*/*JPX* in the respective species, reveal the effect of the environment on LRG mode of action but do not interrogate LRGs’ function in their endogenous contexts. The lack of *XIST* deregulation upon *JPX* RNA depletion in all human cellular contexts that we tested, together with the fact that deleting *JPX* impacts on *XIST* only in *cis* strongly argues against a major role for *JPX* RNA in *trans* during human XCI.

The mechanistic diversification we observed between mouse and human might be a consequence of the changes within the chromatin neighborhood encompassing the *JPX* locus in human. For instance, our previous work suggested that transcriptional activity of *Xist* in the mouse is, at least partially, regulated in *cis* by transcription of the neighboring *Ftx* LRG, independently of the *Ftx* RNA products (Furlan et al., 2018). Interestingly, *Ftx* is located 141 kb upstream of *Xist*, which is comparable to the distance bridging *XIST* promoter to the interaction hotspot Ht1 (∼163 kb) within the human *JPX*, and interacts through CTCF-mediated loops with *Xist* (Furlan et al., 2018). It is within this insulated chromatin neighborhood, which has been reshaped between mouse and human, that constraints on the *JPX* locus might have favored diversification of *JPX* mode of action on *XIST*. One compelling hypothesis from this model is that *XIST* transcriptional *cis*-regulators in eutherian species could have been co-opted based on features such as linear distance from *XIST* promoter, local 3D organization and XCI escaping profile. Importantly, this scenario is reminiscent of what has been observed for enhancer evolution (Villar et al., 2015), and is thus likely not restricted to *JPX* evolution but may apply to other orthologous LRGs.

Finally, what our study provides is a proof of concept that orthologues may act differently in various species, thus epitomizing the mechanistic plasticity of LRGs through evolution. Diversification of LRGs across evolution could confer molecular drift to developmental processes, contribute to species adaptability and fitness: their strong turnover offers a plausible mechanism for generating phenotypic diversity in the control of gene expression across evolution. One major challenge is the systematic identification of versatile LRGs. Indeed, both LRGs functionality, if any, and their mechanism of action are hardly predictable based on the DNA sequence alone. Like for *JPX*, syntenic LRGs often display strong primary sequence turnover during evolution, even among closely related species (Hezroni et al., 2015; Necsulea et al., 2014; Ulitsky et al., 2011; Washietl et al., 2014). The impact of such turnover on LRGs functional conservation is still poorly understood. In rare studies where the functional conservation of lncRNA molecules has been addressed, orthologues display short patches of conserved sequence that are necessary, but not sufficient, for their function (Lin et al., 2014; Ulitsky et al., 2011). This contrasts sharply with the evolutionary stability of protein-coding genes and has often raised controversies about LRGs functionality. Our study pave the way for systematic experimental investigations of LRGs functional conservation with the aim to provide a definitive understanding of underlying rules. Whether mechanistic or functional, this plasticity appears to be an essential parameter to take into consideration in the context of animal modelling of human diseases involving LRGs.

## Data and materials availability

All data generated or analyzed during this study are included in the published article (and its supplementary information).

## Acknowledgments

We thank lab members for critical evaluation of the work leading to this publication. We also thank Antonin Morillon, Pierre-Antoine Defossez, Claire Francastel, Jonathan Weitzman and Céline Morey for critical reading of the manuscript. We thank the Epigenomic, the Microscopy, the Vectorology and the Bioinformatics Platforms, all hosted in UMR7216 Epigenetic and Cell Fate, for technical advices and access to instruments. We acknowledge the ImagoSeine core facility of the Institut Jacques Monod, member of the France BioImaging (ANR-10-INBS-04) and the support of the Region Île-de-France (E539). The research leading to these results has received funding from the European Commission Network of Excellence EpiGeneSys (HEALTH-F4-2010-257082, to C.R. and P.J.R.-G.), from the Agence Nationale pour la Recherche (ANR-14-CE10-0017, to C.R.) and from the Ligue Nationale contre le Cancer (to C.R.). This study was supported by the LabEx “Who Am I?” (ANR-11-LABX-0071) and the Université de Paris IdEx (ANR-18-IDEX-0001) funded by the French Government through its “Investments for the Future” program. O.R. is supported by fellowships from the French Ministry of Education and Research and from the French Medical Research Foundation (FRM). P.J.R.-G. and A.J.C. are supported by the Biotechnology and Biological Sciences Research Council (BB/M022285/1 and BB/P013406/1) and the Medical Research Council (MR/J003808/1).

## Author contributions

O.R., J.-F.O. and C.R. conceived the project and planned the experiments. O.R., C.H., M.C. and J.-F.O. performed the experiments. A.J.C. and P.J.R.-G. performed LGs experiment on naïve hESCs. O.R., J.-F.O. and C.R wrote the manuscript. All authors commented on and revised the manuscript.

## Materials & Correspondence

correspondence and material requests should be addressed to C.R and J.-F.O.

## STAR METHODS

### Cell culture

Primary mouse embryonic fibroblasts (pMEFs) and primary fetal lung fibroblast (IMR90, ATCC CCL-186) were cultured in Dulbecco′s modified Eagle medium (DMEM, Gibco) supplemented with 10% of heat-inactivated Fetal Bovine Serum (FBS, Gibco), 100U/mL of penicillin and 100µg/mL of streptomycin (Thermo Fisher Scientific). Cells were routinely passaged 0.05% Trypsin-EDTA (Thermo Fisher Scientific) and cultured in 20% O2 and 8% CO2 at 37°C.

Female EpiSCs (gift from Alice Jouneau) were cultured using chemically defined medium (CDM) as previously defined(Brons et al., 2007), supplemented with Activin A (20ng/mL, Cell Guidance System) and Fgf2 (12 ng/mL, Cell Guidance System). EpiSCs were passaged using 4 mg/mL Collagenase II (Sigma) and then plated into plates pre-coated with fetal bovine serum.

H9(Thomson et al., 1998) and WIBR2 (Lengner et al., 2010) primed ES cells were cultured on Matrigel-coated culture dishes (BD Biosciences) in mTeSR™1 media (Stemcells technologies) according to the manufacturer instructions, in 20% O2 and 5% CO2 at 37°C. Primed hESCs were routinely passaged in clumps using a 0.5mM EDTA solution as previously described(Beers et al., 2012). For experiments requiring single-cell suspension, cells were incubated with Accutase (Stemcells technologies) and plated in fresh mTeSR™1 media supplemented with 10µM of Y-27632 (Stemcells technologies).

Naïve H9 NK2 cells(Takashima et al., 2014) were cultured in 5% O2 and 5% CO2 at 37°C on CF1 MEFs in a 1:1 mixture of DMEM-F12 and Neurobasal (Thermo Fisher Scientific), 0.5x N2-supplement (Thermo Fisher Scientific), 0.5x B27-supplement (Thermo Fisher Scientific), 1x Non-Essential Amino Acids (Thermo Fisher Scientific), 2mM L-Glutamine (Thermo Fisher Scientific), 1x Penicillin/Streptomycin (Thermo Fisher Scientific), 0.1mM β-mercaptoethanol (Sigma-Aldrich), 1µM PD0325901, 1µM CHIR99021, 20ng/ml human LIF (all from WT-MRC Cambridge Stem Cell Institute) and 2µM Gö6983 (Tocris).

### Derivation of primary MEFs

pMEFs were derived from 13.5 days post-coitum embryos obtained from crosses of CD1 and Crl:CD1(ICR) mice (Charles River). Embryos were manually cut, further dissociated in 0.05% Trypsin-EDTA (Thermo Fisher Scientific) and plated on gelatin-coated dishes. At confluence, pMEFs were frozen until further use (passage 1) and all experiments were performed between passage 1 and 4, from at least three independent female embryos.

### Conversion of primed to naïve hESCs

Prior to conversion, primed hESCs were cultured with mTeSR™1 media (see above) in hypoxia (5% O2 and 5% CO2) for three passages. Primed hESCs were converted into a naïve-like state using the NaïveCult™ Induction Kit cells (STEMCELL technologies) following the guidelines of the manufacturer. Briefly, resetting was launched using 200 000 single-cells plated onto a layer of immortalized MEFs under hypoxic condition. Cells were routinely passaged using TrypLE express enzyme (Thermo Fisher Scientific). RNA samples and cells for RNA-FISH analysis for primed and naïve hESCs were collected at different passages in mTeSR™1 in hypoxia and in expansion medium (day 43/passage 6) respectively. RT-qPCR primers were previously described (Wang et al., 2014).

### LNA GapmRs and si-RNAs lipofection

All LNA Gapmers (LGs) were designed using the Exiqon online tool (https://www.exiqon.com/) and the siRNA targeting YY1 was previously described(Makhlouf et al., 2014). A non-targeting LG and si-RNA were used as negative controls. LGs were lipofected using the RNAi Max transfection reagent (Invitrogen) according to the manufacturer recommendations. Except for Naïve hESCs, all LNA GapmRs (LGs) experiments were performed at a final concentration of 50nM (30 nM siRNAs) using a reverse transfection protocol. For naïve H9-NK2 hESCs, cells were lipofected twice first at 0h by reverse transfection and at 24h by forward transfection, with a final concentration of 25 nM of LGs. All samples were collected 48h post-lipofection either in TRIzol for RNA extraction or Laemmli for western blot analysis. The LGs sequences are listed below:

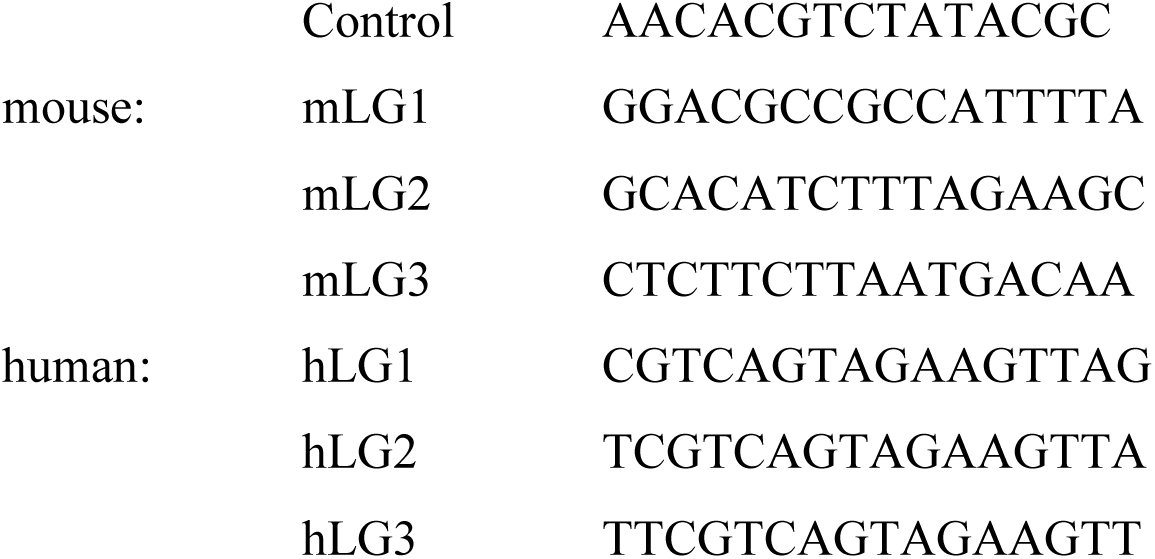

### Total RNA extraction and RT-qPCR

Total RNAs were collected using TRIzol (Thermo Fisher Scientific) and extracted following the manufacturer’s instruction. RNA Samples were treated using the DNA free Kit (Thermo Fisher Scientific) following the manufacturer recommendations. RNAs were reverse transcribed for 30 min at 50°C using the Superscript IV kit (Thermo Fisher Scientific). cDNAs were diluted 1:5 in water and transcripts expression level was assessed by real-time quantitative PCR (RT-qPCR) using the Power SYBR Green Master Mix (Thermo Fisher Scientific). All samples were run in duplicate on a ViiA-7 real-time thermal cycler (Applied Biosystems). Transcripts RNA levels were normalized against a reference gene following the 2-ΔCt method. Unless stated, the Rplp0 gene was used as a reference mouse samples and GAPDH for human samples. All the RT-qPCR primers used in this study are listed in the Table S1.

### Western Blot

Total proteins were extracted with Laemmli lysis buffer (4% SDS; 20% glycerol; 10% 2-mercaptoethanol; 0.004% bromophenol blue; 0.125M Tris-HCl) and sonicated on a Bioruptor Sonication System (Diagenode, UCD-200). After 5 min denaturation at 95°C, the samples were loaded into a 4-12% gradient polyacrylamide gel (Invitrogen) for SDS-PAGE electrophoresis and transferred onto Invitrolon PVDF membranes (Invitrogen). The membranes were blocked for 1h with 5% milk in TBST (10 mM Tris, pH 8.0, 150 mM NaCl, 0.5% Tween 20) and incubated overnight at 4 °C with antibodies targeting YY1 (1:500, mouse sc-7341, H-10, Santa Cruz Biotechnology) or VINCULIN (1:2000, mouse V9131, Sigma Aldrich) proteins. Proteins of interest were detected using a Peroxidase-conjugated antibody (Goat anti-mouse, 1:10 000, Sigma Aldrich) with the Pierce ECL Western blotting substrate (Thermo Scientific).

### Nascent RNA pulldown

Nascent RNAs were purified using the Click-iT Nascent RNA capture kit (Invitrogen). Briefly, cells were incubated for 1h at 37°C with DMEM supplemented with 0.5 mM final of Ethynyl Uridine and total RNAs were extracted using TRIzol reagent. 2µg of total RNAs were used for the biotinylation reaction using 0.5 mM of biotin azide. 1µg of biotinylated RNAs were used for the pulldown assay using Dynabeads MyOne Streptavidin T1 magnetic beads. Reverse transcription was performed using the Superscript VILO cDNA Synthesis Kit (Invitrogen) for 1h at 42°C. cDNAs were diluted 1:2 before RT-qPCR to quantify nascent gene expression. For each sample, a condition without EU was processed in parallel (EU-) and a 10% input (biotinylated RNA before IP) was used to assess enrichment of EU-labelled transcripts after pulldown, in both EU+ and EU-conditions. The values presented in the figures represents Δ(−Ct) values levels normalized to the nascent level of the H2A gene.

### RNA-FISH

Cells preparation. Naïve H9-NK2 and primed hESCs were grown on coverslips. pMEFs, IMR90 and naïve H9 lines were centrifuged onto SuperfrostPlus slides (VWR) using the Cytospin 3 Cytocentrifuge (Shandon). The cells were fixed for 10 min in a 3% Paraformaldehyde solution (Electron Microscopy Science) and permeabilized for 5-10 min in ice-cold CSK buffer (10mM PIPES; 300mM sucrose; 100mM NaCl; 3mM MgCl2; pH6.8) supplemented with 0.5% Triton X-100 (Sigma-Aldrich), 2mM EGTA (Sigma-Aldrich) and 2mM VRC (New England Biolabs).

Probes preparation. RNA-FISH probes were obtained after Nick translation of fosmids/BAC constructs purified using the Large Construct kit (Qiagen): 1µg of purified DNA was labelled for 3h at 15°C with fluorescent dUTPs (SpectrumOrange and SpectrumGreen from Abott Molecular and Cy5-UTPs from GE HealthCare Life Science). The templates used in this study are listed below:

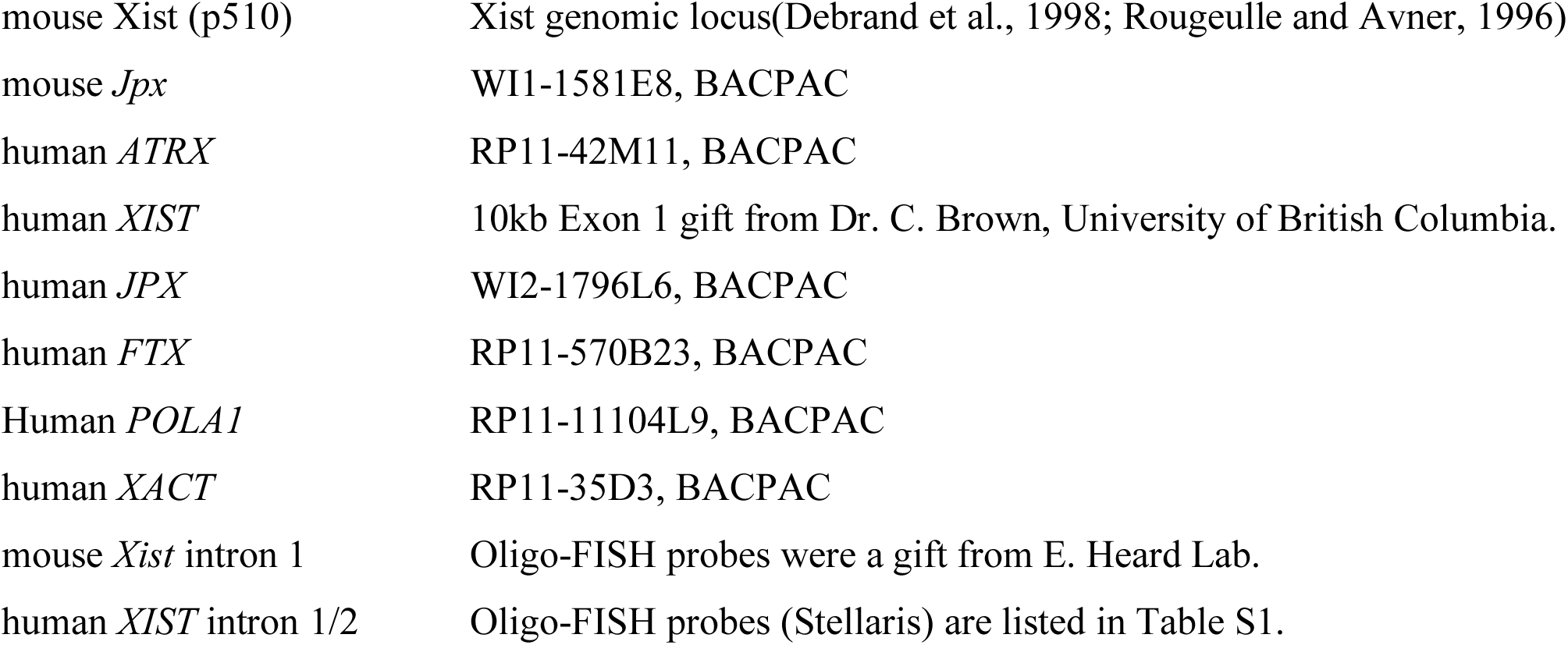

Hybridization. 100 ng of probes were supplemented with 1µg of Cot-I DNA (Invitrogen) and/or 3µg of Sheared Salmon Sperm DNA (Invitrogen). After precipitation, the probes were resuspended in deionized formamide (Sigma Aldrich), denatured for 7 min at 75 °C and further incubated for 15 min at 37 °C if Cot-I DNA was used. Probes were mixed with an equal volume of 2X Hybridization Buffer (4XSSC, 20% Dextran Sulfate, 2mg/ml BSA, 2mM VRC). Coverslips were dehydrated in 80-100% ethanol washes and incubated with the hybridization mix at 37°C overnight in a humid chamber. Next, the coverslips were washed for 4 min at 42°C three times with 50%formaldehyde/2X-SSC (pH7.2) and three times with 2X-SSC. The coverslips were mounted in Vectashield plus DAPI (Vector Laboratories).

### Immunofluorescence coupled to RNA-FISH

Immunofluorescence coupled to RNA-FISH was performed as described previously(Vallot et al., 2017). The antibodies used for IF are listed below:

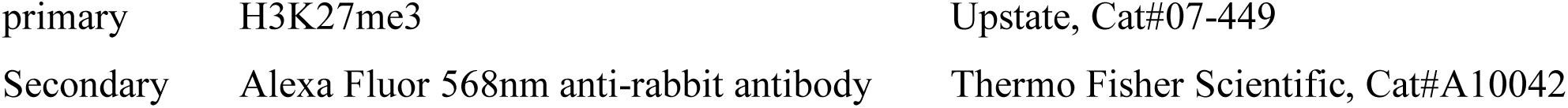

|  |  |  |
| --- | --- | --- |
| primary | H3K27me3 | Upstate, Cat#07-449 |
| Secondary | Alexa Fluor 568nm anti-rabbit antibody | Thermo Fisher Scientific, Cat#A10042 |

### Microscopy and image analysis

All fluorescent microscopy images were taken on a fluorescence DMI-6000 inverted microscope with a motorized stage (Leica), equipped with a CCD Camera HQ2 (Roper Scientifics) and a HCX PL APO 100X oil objective (numerical aperture, 1.4, Leica) using the Metamorph software (version 7.04, Roper Scientifics). Depending on the cell line, 30-60 optical z-sections were collected at 0.2, 0.25 or 0.3µm steps, at different wavelengths depending on the signal (DAPI [360nm, 470nm], FITC [470nm, 525nm], Cy3 [550nm, 570nm], Texas Red [596nm, 612nm] and Cy5 [647nm, 668nm]). Stacks were processed using ImageJ 1.48(Abramoff et al., 2004), and are represented as a 2D “maximum projection” throughout the manuscript. The volume of *XIST* RNA clouds was assessed on stacks using the plugin 3D object counter from ImageJ(Bolte and Cordelieres, 2006).

### Cellular fractionation

Cellular fractionation was performed on at least 5 millions of cells to allow precise estimation of the cells (V) and nuclei (V’) volumes. Fresh pellets of cells were resuspended in 3 volumes (V) of hypotonic buffer (20mM HEPES pH7; 10mM KCl; 0.15mM EDTA; 0.15mM EGTA; 0.15mM spermidine; 0.15mM spermine). Lysis was performed by adding NP-40 (1% final, IGEPAL CA-630) and was stopped with the addition of 0.9V of SR buffer (50mM HEPES pH 7; 0.25mM EDTA; 10mM KCl; 70% sucrose; 0.15mM spermidine; 0.15mM spermine). The cytosolic fraction was separated from the nuclei by 5 min centrifugation at 4°C, 2000g and collected in TRIzol. The pellet of nuclei was washed in 3V of nuclei wash buffer (10mM HEPES pH8; 0.1mM EDTA; 100mM NaCl; 25% glycerol; 0.15mM spermidine; 0.15mM spermine) to remove cytoplasmic contaminations. The volume of the nuclei pellet was estimated (V’) and the nuclei were resuspended in one V’ of sucrose buffer (20mM TRIS pH7.65; 60mM NaCl; 15mM KCl; 0.34M sucrose; 0.15mM spermidine; 0.15mM spermine). The nuclei were incubated for 30 min at 4°C with 0.29V’ of high salt buffer (900mM NaCl; 20mM TRIS pH7.65; 25% glycerol; 1.5mM MgCl2; 0.2mM EDTA) to empty the nuclei of their soluble content. After 30 min centrifugation at 4°C/10 000g, the supernatant and the pellet were collected separately in TRIzol, representing respectively the soluble and non-soluble nuclear fractions. After RT-qPCR, the absolute abundance of the transcripts (Δ-Ct) was normalized to the RNA quantity present in each fraction, from which we computed the abundance of the transcript in a given fraction.

### Chromatin immunoprecipitation

ChIP experiments were performed as described previously(Navarro et al., 2010).

Cells were crosslinked in 1% Formaldehyde (Cliniscience) for 10 min and quenched with 0.125 mM glycine for 5 min. Nuclei were extracted after 30 min incubation in Swelling Buffer (5mM PIPES pH8.0; 85mM KCl; 0.5% NP-40). Samples were then sonicated in TSE150 buffer (0.1% SDS; 1% Triton; 2mM EDTA; 20mM Tris-HCl pH8; 150mM NaCl) using a Bioruptor Sonication System (Diagenode, UCD-200). 1-2µg of antibody were incubated overnight with 5-20µg of chromatin and protein A Magnetic Beads (Thermo Scientific). The following mix was then washed in TSE150, TSE500 (20mM Tris-HCl pH8; 2mM EDTA; 0.1% SDS; 1% Triton X-100; 500mM NaCl), Washing buffer (10mM Tris-HCl pH8; 1mM EDTA; 250mM LiCl; 0.5% NP-40; 0.5% Na-deoxycholate), twice in TE (10mM Tris-HCl pH8; 1mM EDTA) and eluted in TE/1% SDS. After reverse-crosslink (overnight, 65°C), the samples were purified using a phenol-chloroform extraction, resuspended in water and further analyzed by qPCR in duplicates on both IP and input DNA. All values were processed following the 2-ΔCt method and normalized to the input. The primers used for qPCR are available in Table S1. The antibodies used in this study are listed below:

|  |  |
| --- | --- |
| CTCF | Millipore, Cat#07-729, Lot: 2452497 |
| YY1 | Abcam, Cat#ab109237, Lot: GR188694-6 |
| H3K9me3 | Diagenode, Cat#pAb-193-050, Lot: A1671-001P |
| Total RNA Pol2 | Active Motif, Cat#91151 |
| RNA Pol2 CTD Ser5P | Active Motif, Cat#91119 |

### Lentivectors production

Lentiviral particles were produced by transient transfection of HEK293T cells using the calcium-phosphate transfection method. The lentiviral constructs of interest were co-transfected with pMD2.G (Addgene #12259) and psPAX2 (Addgene #12260) plasmids (kindly provided by Didier Trono). After 48h, the culture media was collected, and lentiviral particles were concentrated by ultracentrifugation. For each construct, we assessed the lentiviral titer by infection of HEK293T with serial dilution (1:3) of the lentivirus into DMEM and FACS analysis.

### CRISPR inhibition

The CRISPR inhibitor system(Gilbert et al., 2013) was used to inhibit *JPX* transcription in primed H9 hESCs. DNA oligonucleotides corresponding to the sgRNAs sequences were obtained with the online software CCTop (https://crispr.cos.uni-heidelberg.de/index.html). Oligonucleotide pairs were annealed to generate short double-stranded DNA fragments with overhangs compatible with ligation into the BsmbI-digested plasmid pLKO5.sgRNA.EFS.tGFP (Addgene #57823).

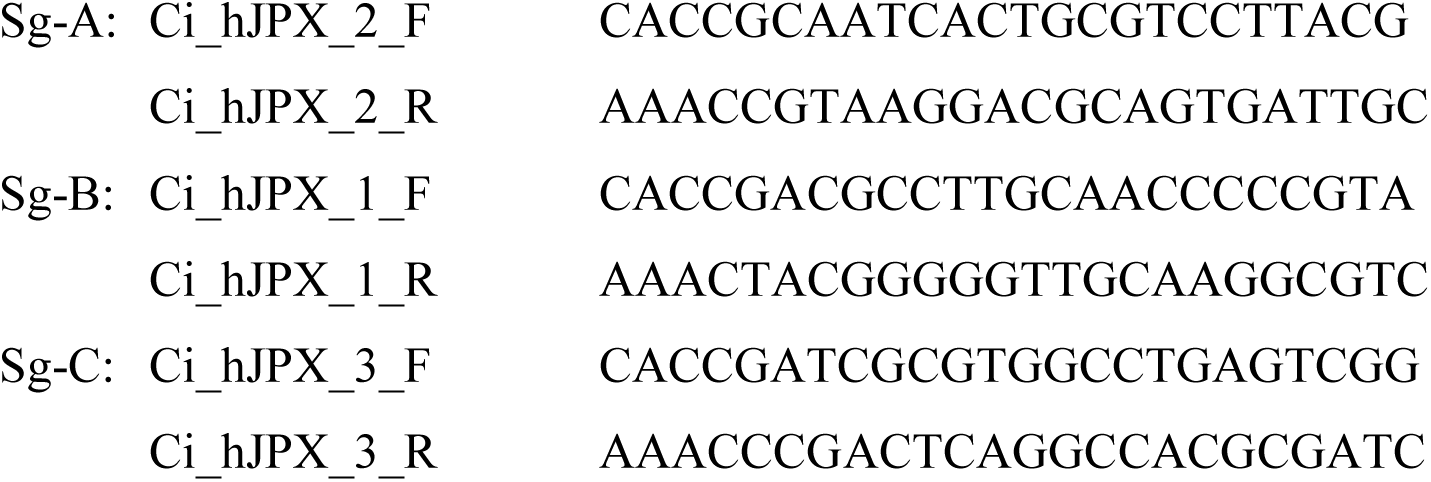

Production of stable cell lines. H9 cells were infected with the dCas9-mCherry-KRAB construct(Furlan et al., 2018) and sorted by FACS (INFLUX 500-BD BioSciences). A second lentiviral infection was performed with the constructs containing the sgRNAs (∼80-85% GFP positive).

### CRISPR/Cas9-mediated deletion of the *JPX* promoter region

*JPX* promoter was deleted in primed H9 hESCs using the CRISPR-Cas9 system. To proceed, plasmid constructs harboring both the sgRNA sequence and the Cas9 fused to a reporter gene were used to allow subsequent selection of transfected cells by FACS. sgRNAs downstream of *JPX* TSS were cloned into a Cas9-GFP construct while upstream guides were cloned into a Cas9-mCherry construct. Therefore, double GFP+/mCherry+ positive cells represent the fraction of cells simultaneously transfected with the two sgRNAs, where the probability for a direct deletion event was increased.

Guides design and cloning. DNA oligonucleotides corresponding to the sgRNAs sequences were obtained with the online software Zifit (http://zifit.partners.org/ZiFiT/ChoiceMenu.aspx).

Oligonucleotide pairs were annealed to generate short double-stranded DNA fragments with overhangs compatible with the ligation into the BbsI-digested plasmid (pSpCas9(BB)-2A-GFP, Addgene #48138, Feng Zhang Lab). We also replaced the GFP by a mCherry reporter to produce a pSpCas9(BB)-2A-mCherry plasmid using the NEBuilder HiFi DNA Assembly Cloning Kit (New England Biolabs). The sequences of the guides are listed below:

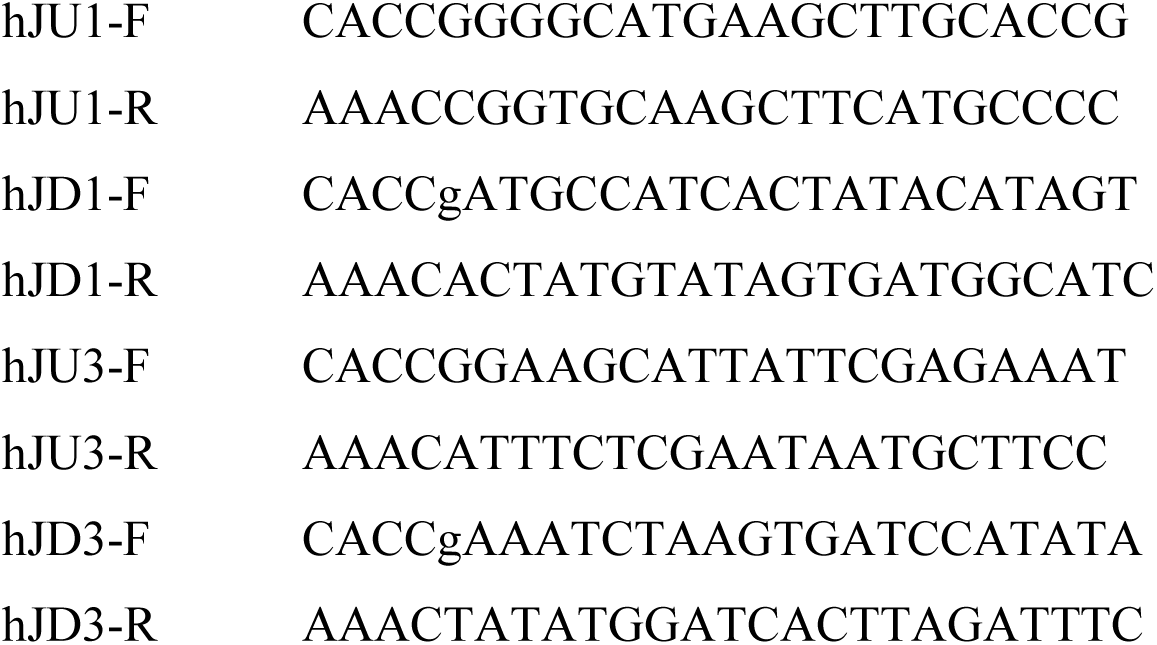

Four million of hESCs were transfected with 5 μg of plasmid DNA for each guide, using the 4D-Nucleofector system (Lonza) as recommended by the manufacturer. 48h post nucleofection, cells were sorted by FACs (INFLUX 500 BD BioSciences) and double positive GFP+ / mCherry+ cells were plated onto Laminin-521 coated plates (Stemcell technologies) at low density in mTeSR™1 supplemented with 1X CloneR™ (Stemcell technologies). Individual colonies were picked and screened by PCR for deletions and inversions events using the following primers:

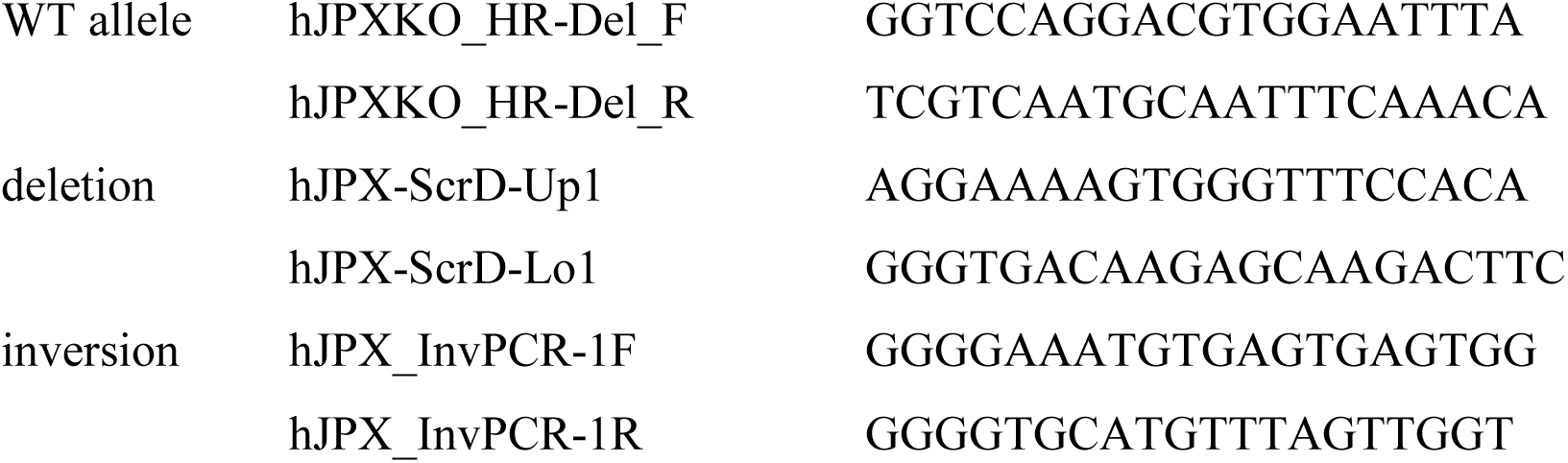

For each clone, the number of X-chromosomes was validated by qPCR on genomic DNA using:

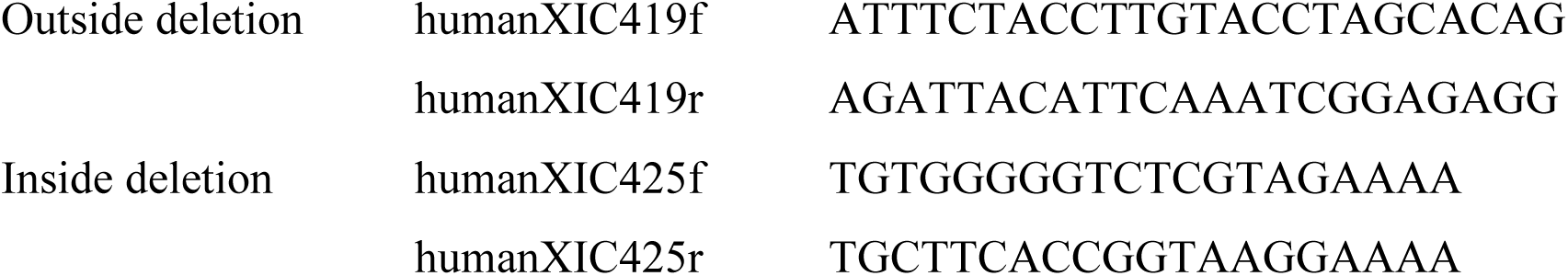

### Resources for genomic data

We downloaded data generated by the ENCODE Project Consortium corresponding to CTCF, RAD21, SMC3, CEPBP, H3K27Ac and H3K4me3 ChIP-seq performed in IMR90 cells; K562 chromatin state hidden Markov model (ChromHMM) and H3K27me3 ChIP-seq and K562. H9 RNA-seq (Vallot et al., 2013) and CTCF ChIP-seq (Ji et al., 2016) were obtained from the GEO repository under the accession numbers GSM978784, GSE62562 and GSE69646 respectively. We obtained sequence conservation of the human *XIC* from the UCSC Genome Browser (Kent et al., 2002) corresponding to the 100 vertebrates Base-wise Conservation by PhyloP.

All heatmap from *in situ* Hi-C datasets (Bonev et al., 2017; Dekker et al., 2017; Rao et al., 2014) represents raw observed matrix visualized at a 5kb resolution and were visualized using the Juicebox suit (Durand et al., 2016a). Domains and loops coordinates were obtained from the corresponding studies.

For ChiA-PET datasets, long-range chromatin interactions and signals tracks were obtained from: (i) the ENCODE project (https://www.encodeproject.org) for POLR2A (ENCSR000BZY) and CTCF (ENCSR000CAC) (Li et al., 2012); (ii) the GEO repository for SMC1 ChIA-PET (GSE69643) (Ji et al., 2016). SMC1 ChIA-PET was processed using Juicer Tools (Durand et al., 2016b) for heatmap visualization. All datas were visualized with the Integrative Genomics Viewer (Robinson et al., 2011) or the UCSC Genome browser(Kent et al., 2002) or on the WashU Epigenome Browser (Zhou et al., 2013; Zhou et al., 2011)).

### Resources for single-cell RNA-Seq

RPKM (Reads Per Kilobase Millions) tables of single-cell RNAseq datasets performed on human embryos (Petropoulos et al., 2016) were obtained from a previous analysis (Vallot et al., 2017). Briefly, RPKM values were computed following a gene-based model and counts falling on regions overlapping two genes were discarded. Unless stated, we used log2(RPKM+0.001) as expression levels for representation and for computation of Pearson’s correlation scores. For lineage assignments, we used the metadata from (Stirparo et al., 2018). All graphical plots were obtained using R (version 3.0.2) with the ggplot2 package (version 1.0.1).

### Statistical information

Throughout the manuscript, RT-qPCR barplots are presented as the mean value with error bars corresponding to standard deviation. The exact number of biological replicates are indicated by the value “n”. Statistical tests used to compute statistical significance are specified in figures legend.

---

**Figure S1:**
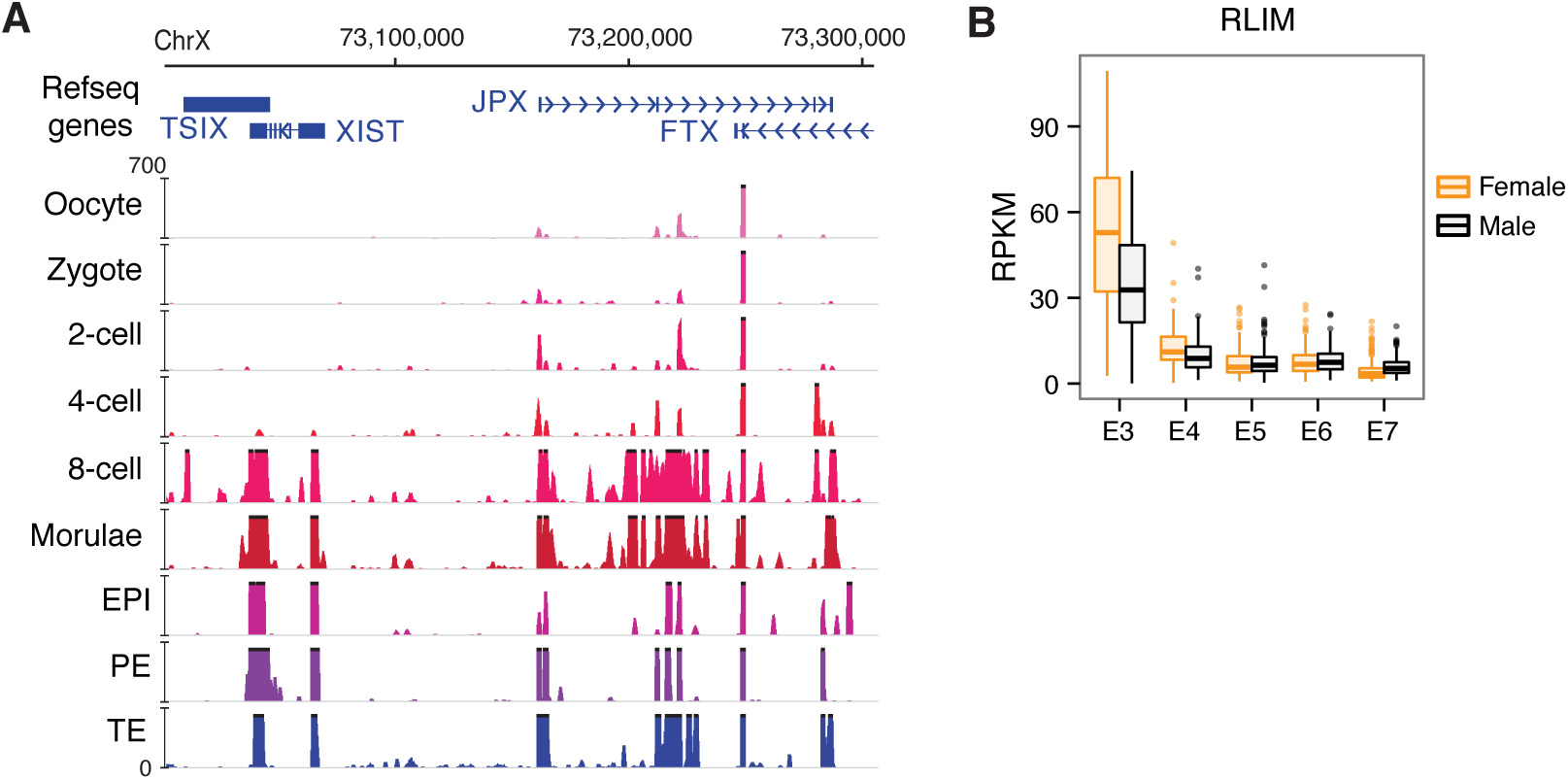
Identification of candidate regulators of *XIST* during early human embryonic development, related to Figure 1. (A) *JPX* reads can be detected from the 2-4 cell stages, possibly linked to limited maternal contribution (Yan et al., 2013), but a burst of *JPX* expression could be seen from the 8-cell stage. At later stages, *JPX* expression can be detected in the three compartments of the embryos – epiblast (EPI), primitive endoderm (PE) and trophectoderm (TE). (B) Expression of the protein-coding gene *RLIM* is the highest at E3 in both male and female embryos and decreases afterwards, suggesting maternal contribution.

**Figure S2:**
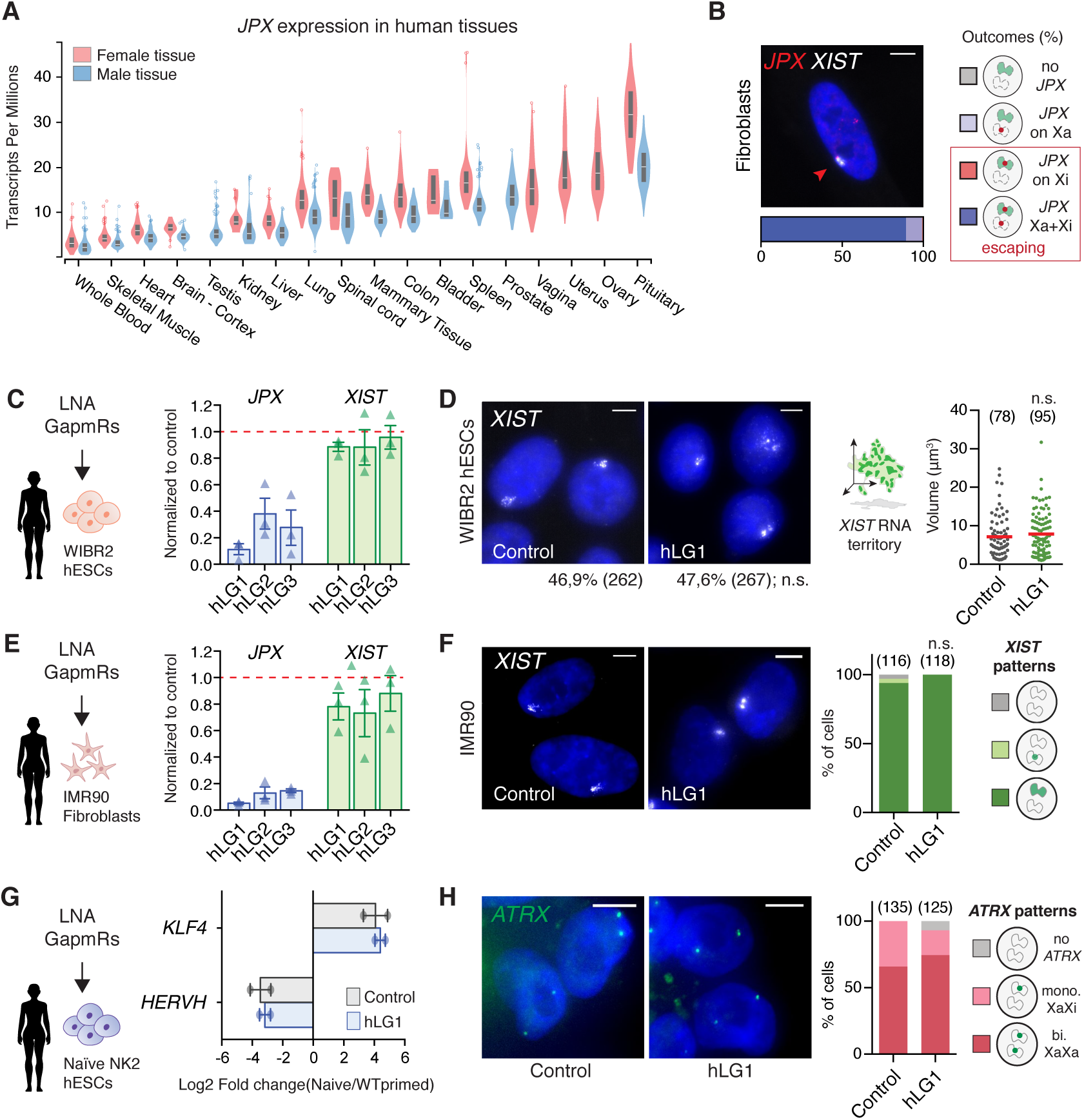
J*P*X RNA is dispensable for *XIST* expression in human, related to Figure 2. (A) *JPX* is ubiquitously expressed across human tissues (Transcripts per Million from the GTEx Project), with ∼2-fold higher expression in female tissues compared to male, in agreement with *JPX* escaping XCI; the box plots shown as median and 25th and 75th percentiles over the violon plots. (B) *JPX* escapes XCI in female fetal fibroblasts, as assessed by *JPX/XIST* double RNA-FISH. (C-D) KD of human *JPX* RNA did not impact *XIST* expression and accumulation in primed WIBR2 hESCs as assessed by RT-qPCR (n=3) and RNA-FISH (Chi square test). (E-F) *XIST* RNA levels were not affected by *JPX* KD in female fetal fibroblasts as assessed by RT-qPCR (n=3) and RNA-FISH (Chi square test). (G) *JPX* KD did not affect the pluripotency status of the cells as assessed by the quantification of naïve (KLF4) and primed (HERVH) markers, RT-qPCR, n=3. (H) *JPX* KD did not affect X-chromosome activity, as determined by *ATRX* RNA-FISH (active X marker). Scale bars are 5 µm. Error bars represent standard deviation; n.s., not significant. Number of counted cells is in brackets.

**Figure S3:**
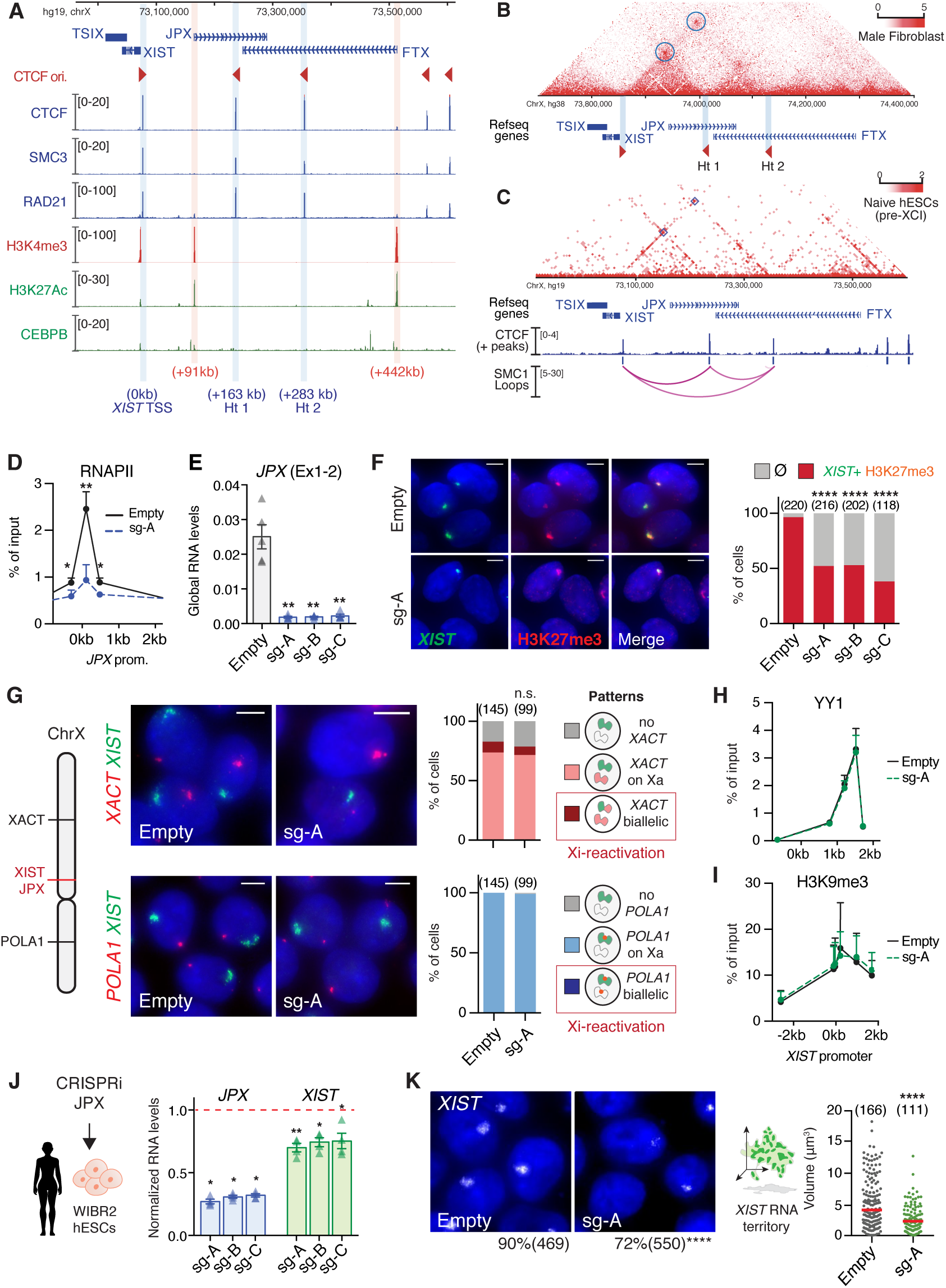
X*I*ST expression requires *JPX* transcription, related to Figure 3. (A) Within the *XIST-*associated TAD, CTCF binding sites are co-occupied by the cohesin complex (RAD21 and SMC3). The hotspots (Ht1/Ht2) are not enriched with enhancer-associated chromatin marks (H3K27Ac) or protein (CEBPB) in female fetal fibroblasts; active promoters in this region are highlighted by the H3K4me3 ChIP-seq track. (B) Structural organization of *XIST-*associated TAD in male fibroblasts (HFF-c6; (Dekker et al., 2017). Called loops are highlighted by blue circles. (C) In naïve hESCs from (Ji et al., 2016), *JPX* and *XIST* are interacting through SMC1-mediated loops, which coincide with CTCF-mediated loops in Fig.2a. The heatmap represents raw SMC1 interactions and arcs represents high confidence loops. (D-E) *JPX* CRISPRi prevented RNAPII recruitment at *JPX* promoter (ChIP-qPCR, n=3) and resulted in a strong decrease of *JPX* RNA levels, RT-qPCR, n=4. (F) *JPX* CRISPRi led to the simultaneous loss of *XIST* RNA clouds (RNA-FISH) and H3K27me3 foci (IF); right panel represents the fraction of double positive cells for *XIST* and H3K27me3 cells, Fischer’s exact test. (G) In contrast to XCI erosion, *JPX* CRISPRi did not trigger *XACT* reactivation from the *XIST*-coated Xi, as assessed by double *XIST* and *XACT* RNA-FISH (Chi-square test). Similarly, no Xi-reactivation of *POLA1* transcription could be identified by double *XIST* and *POLA1* RNA-FISH (Fischer’s exact test). (H) *JPX* CRISPRi did not affect YY1 binding at *XIST* promoter, ChIP-qPCR, n=3. (I) *JPX* CRISPRi did not result in ectopic H3K9me3 enrichment at *XIST* promoter, ChIP-qPCR, n=4. (J-K) Inhibition of *JPX* transcription in WIBR2 hESC induced a decrease in *XIST* RNA levels (RT-qPCR, unpaired two-tailed t-test, n=3), in the number of cells expressing *XIST* (Fischer’s exact test) and in the volume of *XIST* RNA cloud (Mann-Whitney test) – red bars: median. Error bars represent standard deviation; n.s., not significant; *p<0.05; **p<0.01; ***p<0.001; ****p<0.0001. Unpaired two-tailed t-tests to the empty condition unless stated otherwise. Number of counted cells is in brackets.

**Figure S4:**
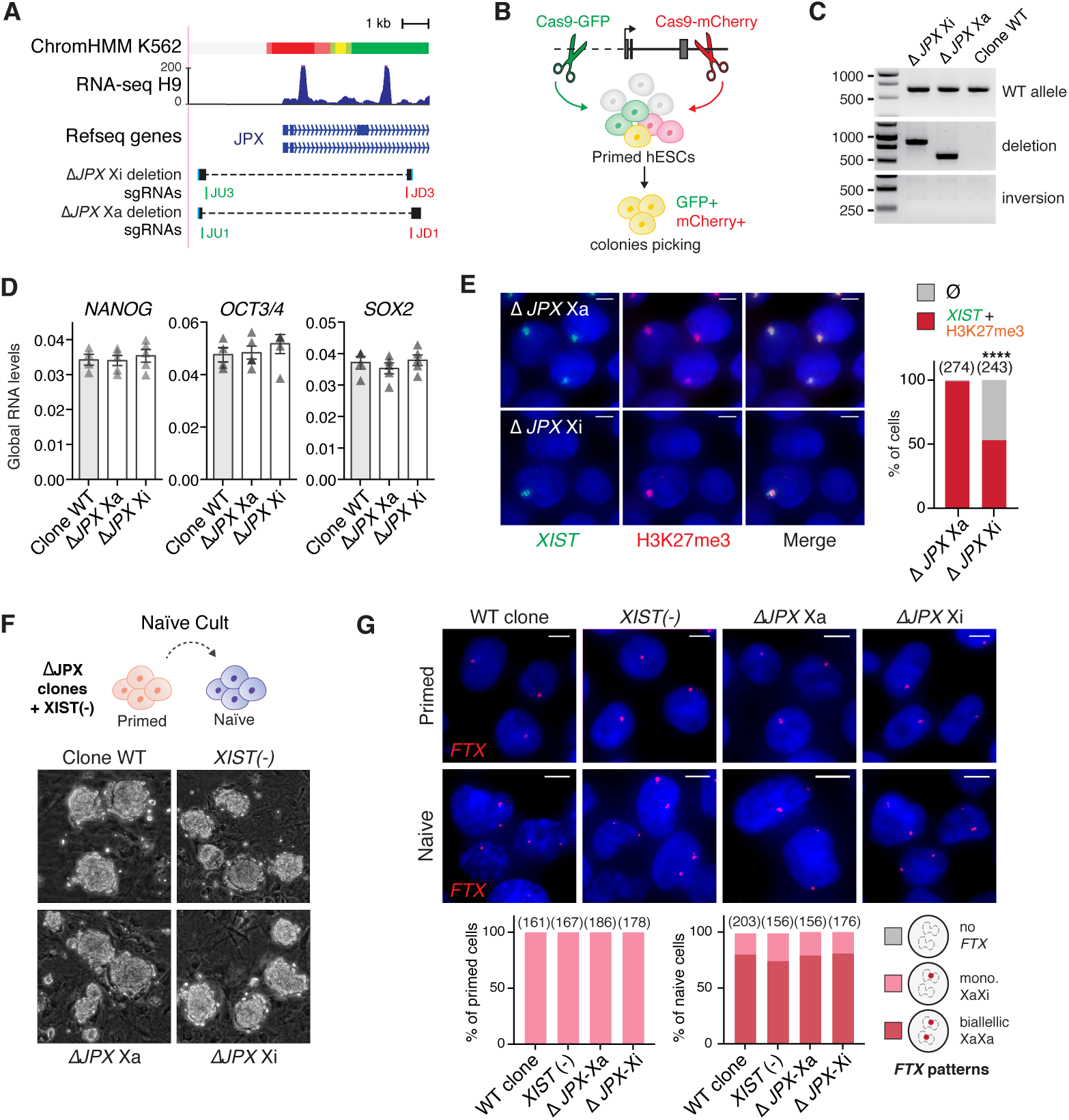
X*I*ST expression requires a functional *JPX* allele in *cis*, related to Figure 4. (A) Map of the *JPX* 5’ region, with various promoter features indicated, such as the chromatin (chromHMM) and transcriptional states (H9 RNA-seq, H3K4me3). The bottom tracks show the guides position (in green, guides coupled to a Cas9-GFP; in red guides coupled to a Cas9-mCherry) and sequence alignments spanning the deleted region to the reference genome in the two clones. (B) Scheme of the strategy to produce genomic deletion using the CRISPR-Cas9 system coupled to FACs-sorting in hESCs. (C) PCR-genotyping of the two heterozygote clones obtained. (D) *JPX* deletion did not impact on the steady state RNA levels of key pluripotency markers, RT-qPCR, error bars represent standard deviation, n=4. (E) Deletion of *JPX* promoter on the Xi, but not the Xa, led to a decrease in the percentage of cells with *XIST* RNA clouds and H3K27me3 foci (Immunofluorescence) (Fischer’s exact test). P-value: <0.0001 (****). (F) Resetting primed hESCs to the naïve state results in similar colonies with dome-shaped appearance, independently of the genotype. (G) Analysis of *FTX* gene expression by RNA-FISH showed robust biallelic expression following resetting of primed into naïve hESCs. Scale bars are 5 µm. Number of counted cells is in brackets.

**Figure S5:**
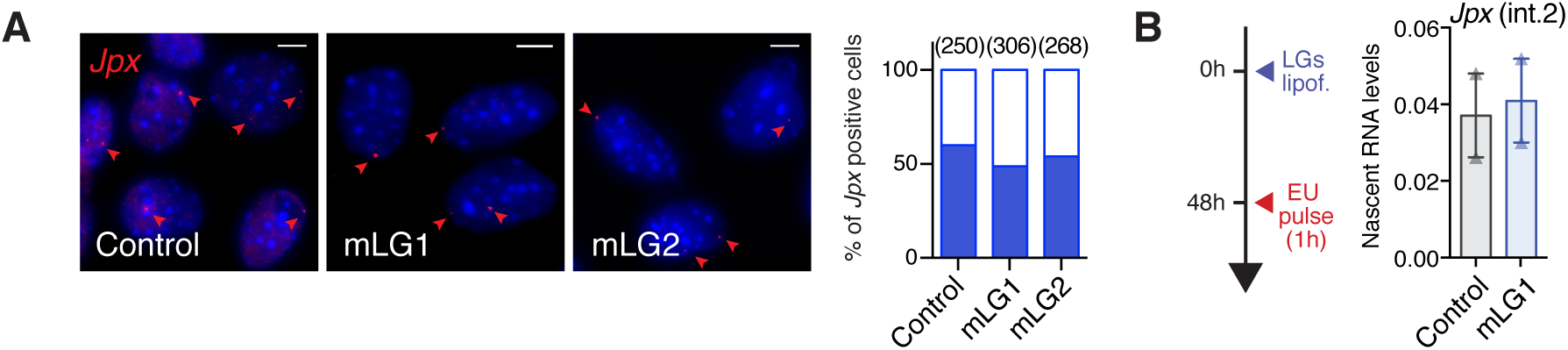
J*p*x RNA regulates *XIST* expression in mouse post-XCI cells, related to Figure 5. (A) RNA-FISH of *Jpx* performed 48h after LGs lipofection in pMEFs. Left panel: representative images. Right panel: quantification of *Jpx* positive cells (blue fill). Number of counted cells is in brackets. (B) Nascent RNA pulldown of EU-labelled nascent transcripts experimental scheme and *Jpx* RNA quantification, RT-qPCR, n=2. Error bars represent standard deviation.

**Figure S6:**
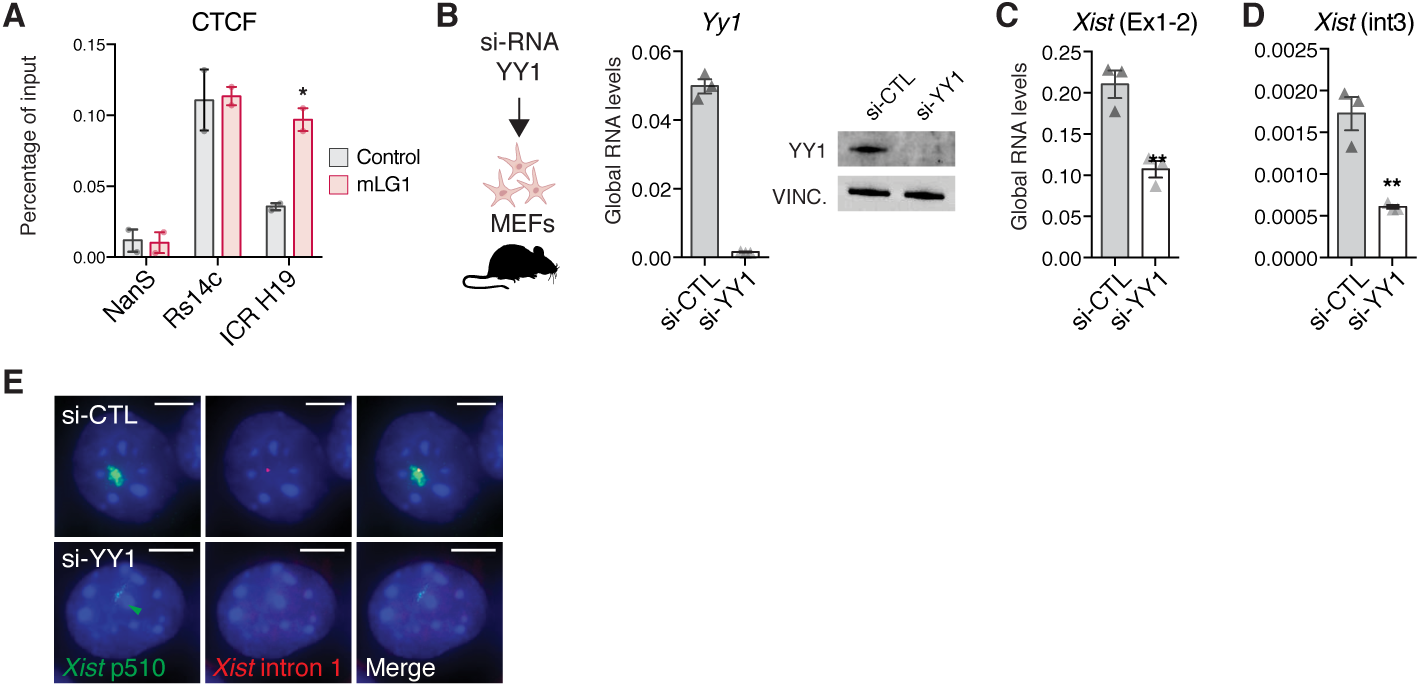
Mechanisms of *XIST* regulation by *JPX* have diversified during evolution, related to Figure 6. (A) CTCF binding at different control positions upon LNA GapmeR transfection. Rs14c and NanS represent respectively positive and negative positions for CTCF binding, ChIP-qPCR, n=3. (B) Expression of *Yy1* mRNA and protein levels following siRNA transfection as in (Makhlouf et al., 2014). (C-D) Expression levels of *Xist* mature and premature RNA following siRNA transfection. (E) Representative images of the effect of *Yy1* depletion on *Xist* accumulation and transcription, RNA-FISH using a probe covering *Xist* locus or intronic probes. Scale bars: 5 µm. Error bars represent standard deviation; unpaired two-tailed t-test, *p<0.05; **p<0.01.

## REFERENCES

Abramoff, M.D., Magalhaes, P.J., and Ram, S.J. (2004). Image Processing with ImageJ. Biophotonics International 11, 36–42.

Beane, R.L., Ram, R., Gabillet, S., Arar, K., Monia, B.P., and Corey, D.R. (2007). Inhibiting gene expression with locked nucleic acids (LNAs) that target chromosomal DNA. Biochemistry 46, 7572–7580.

Beers, J., Gulbranson, D.R., George, N., Siniscalchi, L.I., Jones, J., Thomson, J.A., and Chen, G. (2012). Passaging and colony expansion of human pluripotent stem cells by enzyme-free dissociation in chemically defined culture conditions. Nat Protoc 7, 2029–2040.

Bolte, S., and Cordelieres, F.P. (2006). A guided tour into subcellular colocalization analysis in light microscopy. J Microsc 224, 213–232.

Bonev, B., Mendelson Cohen, N., Szabo, Q., Fritsch, L., Papadopoulos, G.L., Lubling, Y., Xu, X., Lv, X., Hugnot, J.P., Tanay, A., et al. (2017). Multiscale 3D Genome Rewiring during Mouse Neural Development. Cell 171, 557–572 e524.

Brons, I.G., Smithers, L.E., Trotter, M.W., Rugg-Gunn, P., Sun, B., Chuva de Sousa Lopes, S.M., Howlett, S.K., Clarkson, A., Ahrlund-Richter, L., Pedersen, R.A., et al. (2007). Derivation of pluripotent epiblast stem cells from mammalian embryos. Nature 448, 191–195.

Carmona, S., Lin, B., Chou, T., Arroyo, K., and Sun, S. (2018). LncRNA Jpx induces Xist expression in mice using both trans and cis mechanisms. PLoS Genet 14, e1007378.

Carrel, L., Cottle, A.A., Goglin, K.C., and Willard, H.F. (1999). A first-generation X-inactivation profile of the human X chromosome. Proc Natl Acad Sci 96, 14440–14444.

Cho, S.W., Xu, J., Sun, R., Mumbach, M.R., Carter, A.C., Chen, Y.G., Yost, K.E., Kim, J., He, J., Nevins, S.A., et al. (2018). Promoter of lncRNA Gene PVT1 Is a Tumor-Suppressor DNA Boundary Element. Cell 173, 1398–1412 e1322.

Chureau, C., Prissette, M., Bourdet, A., Barbe, V., Cattolico, L., Jones, L., Eggen, A., Avner, P., and Duret, L. (2002). Comparative sequence analysis of the X-inactivation center region in mouse, human, and bovine. Genome Res 12, 894–908.

Debrand, E., Heard, E., and Avner, P. (1998). Cloning and localization of the murine *Xpct* gene: evidence for complex rearrangements during the evolution of the region around the Xist gene. Genomics 48, 296–303.

Dekker, J., Belmont, A.S., Guttman, M., Leshyk, V.O., Lis, J.T., Lomvardas, S., Mirny, L.A., O’Shea, C.C., Park, P.J., Ren, B., et al. (2017). The 4D nucleome project. Nature 549, 219–226.

Dowen, J.M., Fan, Z.P., Hnisz, D., Ren, G., Abraham, B.J., Zhang, L.N., Weintraub, A.S., Schujiers, J., Lee, T.I., Zhao, K., et al. (2014). Control of cell identity genes occurs in insulated neighborhoods in mammalian chromosomes. Cell 159, 374–387.

Durand, N.C., Robinson, J.T., Shamim, M.S., Machol, I., Mesirov, J.P., Lander, E.S., and Aiden, E.L. (2016a). Juicebox Provides a Visualization System for Hi-C Contact Maps with Unlimited Zoom. Cell Syst 3, 99–101.

Durand, N.C., Shamim, M.S., Machol, I., Rao, S.S., Huntley, M.H., Lander, E.S., and Aiden, E.L. (2016b). Juicer Provides a One-Click System for Analyzing Loop-Resolution Hi-C Experiments. Cell Syst 3, 95–98.

Duret, L., Chureau, C., Samain, S., Weissenbach, J., and Avner, P. (2006). The Xist RNA gene evolved in eutherians by pseudogenization of a protein-coding gene. Science 312, 1653–1655.

Elisaphenko, E.A., Kolesnikov, N.N., Shevchenko, A.I., Rogozin, I.B., Nesterova, T.B., Brockdorff, N., and Zakian, S.M. (2008). A dual origin of the Xist gene from a protein-coding gene and a set of transposable elements. PLoS One 3, e2521.

Engreitz, J.M., Haines, J.E., Perez, E.M., Munson, G., Chen, J., Kane, M., McDonel, P.E., Guttman, M., and Lander, E.S. (2016). Local regulation of gene expression by lncRNA promoters, transcription and splicing. Nature 539, 452–455.

Furlan, G., Gutierrez Hernandez, N., Huret, C., Galupa, R., van Bemmel, J.G., Romito, A., Heard, E., Morey, C., and Rougeulle, C. (2018). The Ftx Noncoding Locus Controls X Chromosome Inactivation Independently of Its RNA Products. Mol Cell 70, 462–472 e468.

Furlan, G., and Rougeulle, C. (2016). Function and evolution of the long noncoding RNA circuitry orchestrating X-chromosome inactivation in mammals. Wiley Interdiscip Rev RNA 7, 702–722.

Gilbert, L.A., Larson, M.H., Morsut, L., Liu, Z., Brar, G.A., Torres, S.E., Stern-Ginossar, N., Brandman, O., Whitehead, E.H., Doudna, J.A., et al. (2013). CRISPR-mediated modular RNA-guided regulation of transcription in eukaryotes. Cell 154, 442–451.

Giorgetti, L., Galupa, R., Nora, E.P., Piolot, T., Lam, F., Dekker, J., Tiana, G., and Heard, E. (2014). Predictive polymer modeling reveals coupled fluctuations in chromosome conformation and transcription. Cell 157, 950–963.

Guo, G., von Meyenn, F., Rostovskaya, M., Clarke, J., Dietmann, S., Baker, D., Sahakyan, A., Myers, S., Bertone, P., Reik, W., et al. (2017). Epigenetic resetting of human pluripotency. Development 144, 2748–2763.

Hezroni, H., Ben-Tov Perry, R., Meir, Z., Housman, G., Lubelsky, Y., and Ulitsky, I. (2017). A subset of conserved mammalian long non-coding RNAs are fossils of ancestral protein-coding genes. Genome Biol 18, 162.

Hezroni, H., Koppstein, D., Schwartz, M.G., Avrutin, A., Bartel, D.P., and Ulitsky, I. (2015). Principles of long noncoding RNA evolution derived from direct comparison of transcriptomes in 17 species. Cell Rep 11, 1110–1122.

Hnisz, D., Day, D.S., and Young, R.A. (2016). Insulated Neighborhoods: Structural and Functional Units of Mammalian Gene Control. Cell 167, 1188–1200.

Ji, X., Dadon, D.B., Powell, B.E., Fan, Z.P., Borges-Rivera, D., Shachar, S., Weintraub, A.S., Hnisz, D., Pegoraro, G., Lee, T.I., et al. (2016). 3D Chromosome Regulatory Landscape of Human Pluripotent Cells. Cell Stem Cell 18, 262–275.

Johnston, C.M., Newall, A.E., Brockdorff, N., and Nesterova, T.B. (2002). Enox, a novel gene that maps 10 kb upstream of Xist and partially escapes X inactivation. Genomics 80, 236–244.

Karner, H.M., Webb, C., Carmona, S., Liu, Y., Lin, B., Erhard, M., Chan, D., Baldi, P., Spitale, R.C., and Sun, S. (2019). Functional conservation of lncRNA JPX despite sequence and structural divergence. bioRxiv 686113.

Kent, W.J., Sugnet, C.W., Furey, T.S., Roskin, K.M., Pringle, T.H., Zahler, A.M., and Haussler, D. (2002). The human genome browser at UCSC. Genome Res 12, 996–1006.

Kolesnikov, N.N., and Elisaphenko, E.A. (2010). Comparative organization and the origin of noncoding regulatory RNA genes from X-chromosome inactivation center of human and mouse. Russian Journal of Genetics 46, 1223–1228.

Lengner, C.J., Gimelbrant, A.A., Erwin, J.A., Cheng, A.W., Guenther, M.G., Welstead, G.G., Alagappan, R., Frampton, G.M., Xu, P., Muffat, J., et al. (2010). Derivation of pre-X inactivation human embryonic stem cells under physiological oxygen concentrations. Cell 141, 872–883.

Leucci, E., Vendramin, R., Spinazzi, M., Laurette, P., Fiers, M., Wouters, J., Radaelli, E., Eyckerman, S., Leonelli, C., Vanderheyden, K., et al. (2016). Melanoma addiction to the long non-coding RNA SAMMSON. Nature 531, 518–522.

Li, G., Ruan, X., Auerbach, R.K., Sandhu, K.S., Zheng, M., Wang, P., Poh, H.M., Goh, Y., Lim, J., Zhang, J., et al. (2012). Extensive promoter-centered chromatin interactions provide a topological basis for transcription regulation. Cell 148, 84–98.

Lin, N., Chang, K.Y., Li, Z., Gates, K., Rana, Z.A., Dang, J., Zhang, D., Han, T., Yang, C.S., Cunningham, T.J., et al. (2014). An evolutionarily conserved long noncoding RNA TUNA controls pluripotency and neural lineage commitment. Mol Cell 53, 1005–1019.

Luo, S., Lu, J.Y., Liu, L., Yin, Y., Chen, C., Han, X., Wu, B., Xu, R., Liu, W., Yan, P., et al. (2016). Divergent lncRNAs Regulate Gene Expression and Lineage Differentiation in Pluripotent Cells. Cell Stem Cell 18, 637–652.

Makhlouf, M., Ouimette, J.F., Oldfield, A., Navarro, P., Neuillet, D., and Rougeulle, C. (2014). A prominent and conserved role for YY1 in Xist transcriptional activation. Nature communications 5, 4878.

Mekhoubad, S., Bock, C., de Boer, A.S., Kiskinis, E., Meissner, A., and Eggan, K. (2012). Erosion of dosage compensation impacts human iPSC disease modeling. Cell Stem Cell 10, 595–609.

Navarro, P., Oldfield, A., Legoupi, J., Festuccia, N., Dubois, A., Attia, M., Schoorlemmer, J., Rougeulle, C., Chambers, I., and Avner, P. (2010). Molecular coupling of Tsix regulation and pluripotency. Nature 468, 457–460.

Navarro, P., Pichard, S., Ciaudo, C., Avner, P., and Rougeulle, C. (2005). Tsix transcription across the Xist gene alters chromatin conformation without affecting Xist transcription: implications for X-chromosome inactivation. Genes Dev 19, 1474–1484.

Necsulea, A., Soumillon, M., Warnefors, M., Liechti, A., Daish, T., Zeller, U., Baker, J.C., Grutzner, F., and Kaessmann, H. (2014). The evolution of lncRNA repertoires and expression patterns in tetrapods. Nature 505, 635–640.

Nora, E.P., Lajoie, B.R., Schulz, E.G., Giorgetti, L., Okamoto, I., Servant, N., Piolot, T., van Berkum, N.L., Meisig, J., Sedat, J., et al. (2012). Spatial partitioning of the regulatory landscape of the X-inactivation centre. Nature 485, 381–385.

Okamoto, I., Patrat, C., Thepot, D., Peynot, N., Fauque, P., Daniel, N., Diabangouaya, P., Wolf, J.P., Renard, J.P., Duranthon, V., et al. (2011). Eutherian mammals use diverse strategies to initiate X-chromosome inactivation during development. Nature 472, 370–374.

Paralkar, V.R., Taborda, C.C., Huang, P., Yao, Y., Kossenkov, A.V., Prasad, R., Luan, J., Davies, J.O., Hughes, J.R., Hardison, R.C., et al. (2016). Unlinking an lncRNA from Its Associated cis Element. Mol Cell 62, 104–110.

Petropoulos, S., Edsgard, D., Reinius, B., Deng, Q., Panula, S.P., Codeluppi, S., Reyes, A.P., Linnarsson, S., Sandberg, R., and Lanner, F. (2016). Single-Cell RNA-Seq Reveals Lineage and X Chromosome Dynamics in Human Preimplantation Embryos. Cell 167, 285.

Rao, S.S., Huntley, M.H., Durand, N.C., Stamenova, E.K., Bochkov, I.D., Robinson, J.T., Sanborn, A.L., Machol, I., Omer, A.D., Lander, E.S., et al. (2014). A 3D map of the human genome at kilobase resolution reveals principles of chromatin looping. Cell 159, 1665–1680.

Romito, A., and Rougeulle, C. (2011). Origin and evolution of the long non-coding genes in the X-inactivation center. Biochimie 93, 1935–1942.

Rougeulle, C., and Avner, P. (1996). Cloning and characterization of a murine brain specific gene *Bpx* and its human homologue lying within the *Xic* candidate region. Hum Mol Genet 5, 41–49.

Sahakyan, A., Kim, R., Chronis, C., Sabri, S., Bonora, G., Theunissen, T.W., Kuoy, E., Langerman, J., Clark, A.T., Jaenisch, R., et al. (2017). Human Naive Pluripotent Stem Cells Model X Chromosome Dampening and X Inactivation. Cell Stem Cell 20, 87–101.

Shin, J., Bossenz, M., Chung, Y., Ma, H., Byron, M., Taniguchi-Ishigaki, N., Zhu, X., Jiao, B., Hall, L.L., Green, M.R., et al. (2010). Maternal Rnf12/RLIM is required for imprinted X-chromosome inactivation in mice. Nature 467, 977–981.

Stirparo, G.G., Boroviak, T., Guo, G., Nichols, J., Smith, A., and Bertone, P. (2018). Integrated analysis of single-cell embryo data yields a unified transcriptome signature for the human pre-implantation epiblast. Development 145.

Stojic, L., Lun, A.T.L., Mangei, J., Mascalchi, P., Quarantotti, V., Barr, A.R., Bakal, C., Marioni, J.C., Gergely, F., and Odom, D.T. (2018). Specificity of RNAi, LNA and CRISPRi as loss-of-function methods in transcriptional analysis. Nucleic Acids Res 46, 5950–5966.

Sun, F., Chronis, C., Kronenberg, M., Chen, X.F., Su, T., Lay, F.D., Plath, K., Kurdistani, S.K., and Carey, M.F. (2019). Promoter-Enhancer Communication Occurs Primarily within Insulated Neighborhoods. Mol Cell 73, 250–263 e255.

Sun, S., Del Rosario, B.C., Szanto, A., Ogawa, Y., Jeon, Y., and Lee, J.T. (2013). Jpx RNA activates Xist by evicting CTCF. Cell 153, 1537–1551.

Takashima, Y., Guo, G., Loos, R., Nichols, J., Ficz, G., Krueger, F., Oxley, D., Santos, F., Clarke, J., Mansfield, W., et al. (2014). Resetting transcription factor control circuitry toward ground-state pluripotency in human. Cell 158, 1254–1269.

Tesar, P.J., Chenoweth, J.G., Brook, F.A., Davies, T.J., Evans, E.P., Mack, D.L., Gardner, R.L., and McKay, R.D. (2007). New cell lines from mouse epiblast share defining features with human embryonic stem cells. Nature 448, 196–199.

Theunissen, T.W., Friedli, M., He, Y., Planet, E., O’Neil, R.C., Markoulaki, S., Pontis, J., Wang, H., Iouranova, A., Imbeault, M., et al. (2016). Molecular Criteria for Defining the Naive Human Pluripotent State. Cell Stem Cell 19, 502–515.

Thomson, J.A., Itskovitz-Eldor, J., Shapiro, S.S., Waknitz, M.A., Swiergiel, J.J., Marshall, V.S., and Jones, J.M. (1998). Embryonic stem cell lines derived from human blastocysts. Science 282, 1145–1147.

Tian, D., Sun, S., and Lee, J.T. (2010). The long noncoding RNA, Jpx, is a molecular switch for X chromosome inactivation. Cell 143, 390–403.

Tripathi, V., Ellis, J.D., Shen, Z., Song, D.Y., Pan, Q., Watt, A.T., Freier, S.M., Bennett, C.F., Sharma, A., Bubulya, P.A., et al. (2010). The nuclear-retained noncoding RNA MALAT1 regulates alternative splicing by modulating SR splicing factor phosphorylation. Mol Cell 39, 925–938.

Ulitsky, I., Shkumatava, A., Jan, C.H., Sive, H., and Bartel, D.P. (2011). Conserved function of lincRNAs in vertebrate embryonic development despite rapid sequence evolution. Cell 147, 1537–1550.

Vallot, C., Huret, C., Lesecque, Y., Resh, A., Oudrhiri, N., Bennaceur-Griscelli, A., Duret, L., and Rougeulle, C. (2013). *XACT*, a long non-coding transcript coating the active X in human pluripotent cells. Nat Genet 45, 239–241.

Vallot, C., Ouimette, J.F., Makhlouf, M., Feraud, O., Pontis, J., Come, J., Martinat, C., Bennaceur-Griscelli, A., Lalande, M., and Rougeulle, C. (2015). Erosion of X Chromosome Inactivation in Human Pluripotent Cells Initiates with XACT Coating and Depends on a Specific Heterochromatin Landscape. Cell Stem Cell 16, 533–546.

Vallot, C., Ouimette, J.F., and Rougeulle, C. (2016). Establishment of X chromosome inactivation and epigenomic features of the inactive X depend on cellular contexts. Bioessays 38, 869–880.

Vallot, C., Patrat, C., Collier, A.J., Huret, C., Casanova, M., Liyakat Ali, T.M., Tosolini, M., Frydman, N., Heard, E., Rugg-Gunn, P.J., et al. (2017). XACT Noncoding RNA Competes with XIST in the Control of X Chromosome Activity during Human Early Development. Cell Stem Cell 20, 102–111.

Villar, D., Berthelot, C., Aldridge, S., Rayner, T.F., Lukk, M., Pignatelli, M., Park, T.J., Deaville, R., Erichsen, J.T., Jasinska, A.J., et al. (2015). Enhancer evolution across 20 mammalian species. Cell 160, 554–566.

Wang, J., Xie, G., Singh, M., Ghanbarian, A.T., Rasko, T., Szvetnik, A., Cai, H., Besser, D., Prigione, A., Fuchs, N.V., et al. (2014). Primate-specific endogenous retrovirus-driven transcription defines naive-like stem cells. Nature 516, 405–409.

Washietl, S., Kellis, M., and Garber, M. (2014). Evolutionary dynamics and tissue specificity of human long noncoding RNAs in six mammals. Genome Res 24, 616–628.

Yan, L., Yang, M., Guo, H., Yang, L., Wu, J., Li, R., Liu, P., Lian, Y., Zheng, X., Yan, J., et al. (2013). Single-cell RNA-Seq profiling of human preimplantation embryos and embryonic stem cells. Nat Struct Mol Biol 20, 1131–1139.

Zhou, X., Lowdon, R.F., Li, D., Lawson, H.A., Madden, P.A., Costello, J.F., and Wang, T. (2013). Exploring long-range genome interactions using the WashU Epigenome Browser. Nat Methods 10, 375–376.

Zhou, X., Maricque, B., Xie, M., Li, D., Sundaram, V., Martin, E.A., Koebbe, B.C., Nielsen, C., Hirst, M., Farnham, P., et al. (2011). The Human Epigenome Browser at Washington University. Nat Methods 8, 989–990.

